# Spatial heterogeneity and organization of tumor mutation burden and immune infiltrates within tumors based on whole slide images correlated with patient survival in bladder cancer

**DOI:** 10.1101/554527

**Authors:** Hongming Xu, Sunho Park, Jean René Clemenceau, Jinhwan Choi, Nathan Radakovich, Sung Hak Lee, Tae Hyun Hwang

## Abstract

High-TMB (TMB-H) could result in an increased number of neoepitopes from somatic mutations expressed by a patient’s own tumor cell which can be recognized and targeted by neighboring tumor-infiltrating lymphocytes (TILs). Deeper understanding of spatial heterogeneity and organization of tumor cells and their neighboring immune infiltrates within tumors could provide new insights into tumor progression and treatment response. Here we developed and applied computational approaches using digital whole slide images (WSIs) to investigate spatial heterogeneity and organization of regions harboring TMB-H tumor cells and TILs within tumors, and its prognostic utility. In experiments using WSIs from The Cancer Genome Atlas bladder cancer (BLCA), our findings show that WSI-based approaches can reliably predict patient-level TMB status and delineate spatial TMB heterogeneity and co-organization with TILs. TMB-H patients with low spatial heterogeneity enriched with high TILs show improved overall survival indicating a prognostic role of spatial TMB and TILs information in BLCA.

## 1 Introduction

Tumor mutational burden (TMB) is a quantitative genomic biomarker that measures the number of mutations within a tumor. TMB level has been shown to be associated with better prognosis and clinical responses to immune-checkpoint inhibitors in various cancer types such as melanoma, lung cancer and bladder cancer [2,18,21,22,45]. Higher TMB levels are correlated with higher levels of neoantigens expressed by a cancer cell, which could help the neighboring tumor-infiltrating lymphocytes (TILs) to recognize and kill them [1]. Various studies including clinical trials reported that patients with TMB high (TMB-H) and/or high density of TILs within tumors had favorable prognosis and response to immunotherapy in many cancer types [2,16,18,21–26,45]. Recent studies showed that spatial heterogeneity and composition of immune cells in the tumor microenvironment could improve our understanding of how immune environment influences patients’ prognosis and response to treatments, including immunotherapy [28, 30–33]. These findings might suggest that detecting regions harboring TMB-H tumor cells and TILs within the tumor microenvironment and analyzing their spatial architecture could provide new insights into the relationship between spatial TMB and TIL co-arrangement and patient’s outcome.

Tissue-based bulk DNA sequencing (e.g. whole exome sequencing (WES), targeted sequencing, etc.) and mRNA sequencing are widely used to assess patient-level TMB status and quantify TILs in tumors, respectively. However, due to the limited tissue availability, high costs and time-consuming procedures, the clinical utility of tissue-based DNA and mRNA sequencing are limited. In addition, these bulk DNA and mRNA sequencing approaches were not designed to take into account spatial intratumor TMB and immune heterogeneity, thus provide potentially biased samples leading to inconsistent testing results [43]. Although the blood-based TMB measurement (i.e. liquid biopsies) has recently become available, this approach poses similar challenges to tissue-based TMB measurements [3]. The development of single-cell DNA and mRNAseq has revealed a spectrum of tumor cell and immune cell heterogeneity in the patient’s tumor, but these approaches do not provide insight into the spatial organization of tumor and immune cell architecture [26,46–48]. Most recently, spatial transcriptome technologies have enabled mapping of the spatial architecture, composition and interactions of various cell types within the tumor, but simultaneously elucidating both DNA (e.g., TMB status) and RNA-level characteristics of cells is still challenging [49–51].

The use of widely available histopathological images poses a promising alternative. Routine histopathological examination is the gold standard for diagnosis, grading, and quantification of TILs for various cancer types in a clinical setting. With the recent development in deep Learning, computational approaches based on whole slide images (WSIs) have been explored to predict genetic characteristics (e.g., mutation status, gene expression, etc.) present within tumor regions in prostate [12], lung [14], colon, stomach [41], and pan-cancer [37, 38].

WSIs are also being widely used to detect tumor-infiltrating lymphocytes (TILs) and its quantification within the tumor by computational analysis. Recently, Saltz *et al*. (2018) proposed to use convolutional neural networks (CNN) to identify TIL in H&E stained WSIs and showed that spatial composition of TILs within tumors correlated with patient’s prognosis across cancers. Corredor *et al*. [35] and Acs *et al*. (2019) [11] developed the algorithms to segment and detect TILs and used spatial composition and co-orgarnization of TILs and cancer cell within tumors linked to cancer recurrence and progosis in non-small cell lung cancer and melanoma, respectively. Most recently, Abduljabbar *et al*. (2020) performed a study integrating multiregion exome and RNA-sequencing (RNA-seq) data with spatial histology to investigate spatial tumor and immune microenvironment in LUAD and showed that lung adenocarcinoma (LUAD) subgroup with immune cold and low neoantigen burden (i.e., low TMB) was significantly correlated with poorer disease free survival [10]. This study demonstrated that the deep learning approach utilizing digital pathology images could provide a deeper understanding of how spatial composition of tumor and immune cells within tumor microenvironment impact tumor evolution and progression.

Given these studies showing that computational approaches and deep learning algorithms utilizing morphological features present in WSIs could reliably predict characteristics of tumor and immune cells and their spatial organization in tumors, we hypothesize that a carefully designed WSI-based computational method could accurately predict TMB status and TILs in given regions within tumors and be used to dissect spatial heterogeneity of TMB and its co-organization with TILs across regions within tumors. Specifically, we hypothesize that the comprehensive understanding of spatial co-occurrence of TILs with neighboring TMB-H or TMB-low (TMB-L) regions from pathology slides could provide a prognostic utility to identify patient subgroups with distinct survival outcome.

In this work, we first develop and evaluate deep-learning based computational pipelines to predict patient and tumor tile-level (i.e., dividing a WSI into small tiles for analysis) TMB status and TILs. We then use the tile-level TMB status to delineate spatial heterogeneity of TMB within WSIs. We perform a joint spatial analysis of regions harboring predicted TMB status and TILs within the tumor and use the spatial heterogeneity and arrangement information to identify patient subgroups (e.g., TMB-H tumor with low spatial TMB heterogeneity enriched with high density of TILs). To the best of our knowledge, this is the first work to interrogate spatial heterogeneity and organization of TMB with TILs within tumors to evaluate its prognostic utility to stratify patients using WSIs.

In experiments with TCGA Urothelial Bladder Carcinoma (BLCA) and Lung Adenocarcinoma (LUAD) cohorts, we first evaluated the performance of our proposed method using WSIs to predict patient-level TMB status against state-of-the-art methods, including deep learning and multiple instance learning methods. The patient-level evaluation performed is mainly because ground truth TMB status is only assigned per patient. Then we applied our proposed model to predict TMB status at the tile level within WSIs for BLCA cohort and applied entropy measurement to evaluate spatial heterogeneity of TMB within WSIs. Finally, we performed a survival analysis of patient subgroups based on spatial TMB and/or TILs information in BLCA cohort. Identification of patient subgroups based on patient-level TMB status and TMB spatial heterogeneity status indicated that incorporating spatial heterogeneity of TMB could lead better patient stratification in BLCA cohort. We further investigated whether incorporating TILs status with spatial TMB status within the tumor could improve patient prognostication. We found that the integration of predicted TMB status and TIL densities within tumor regions could lead significant better patient risk stratification in BLCA.

## 2 Results

### Whole Slide Image Analysis Workflow

We developed a computational pipeline using WSIs to predict patient-level TMB status and delineate spatial heterogeneity of TMB present in tumors. We also trained a deep learning model to detect TILs and quantify its densities within tumor regions. The aim of our approach is to incorporate spatial TMB heterogeneity with patient-level TMB status and TIL densities to identify patient subgroups that could lead to better patient stratification. The computational analysis workflow is shown in Fig. 1(a), which includes two main modules: automatic TMB prediction and TILs detection. In the automatic TMB prediction module, the trained convolutional neural network (CNN) based tumor detector is first applied to identify tumor regions in the WSI (see Fig. 1(b)). Affinity propagation (AP) clustering is then applied to select a subset of representative tumor regions (see Fig. 1(c)(d)). After that, transfer learning using Xception model is used to convert representative tumor patches into feature vectors. Finally, SVM with linear or RBF kernel is trained and tested on integrated patient-level feature vectors. In the automatic TIL detection module, the trained tile-level Resnet18 deep learning model is utilized to identity TIL regions in the WSI. The ratio of identified TILs pixels over the total number of tumor pixels is quantified as a variable to characterize TIL density inside tumor regions. More technical details can be referred in the method section.

**Figure 1:**
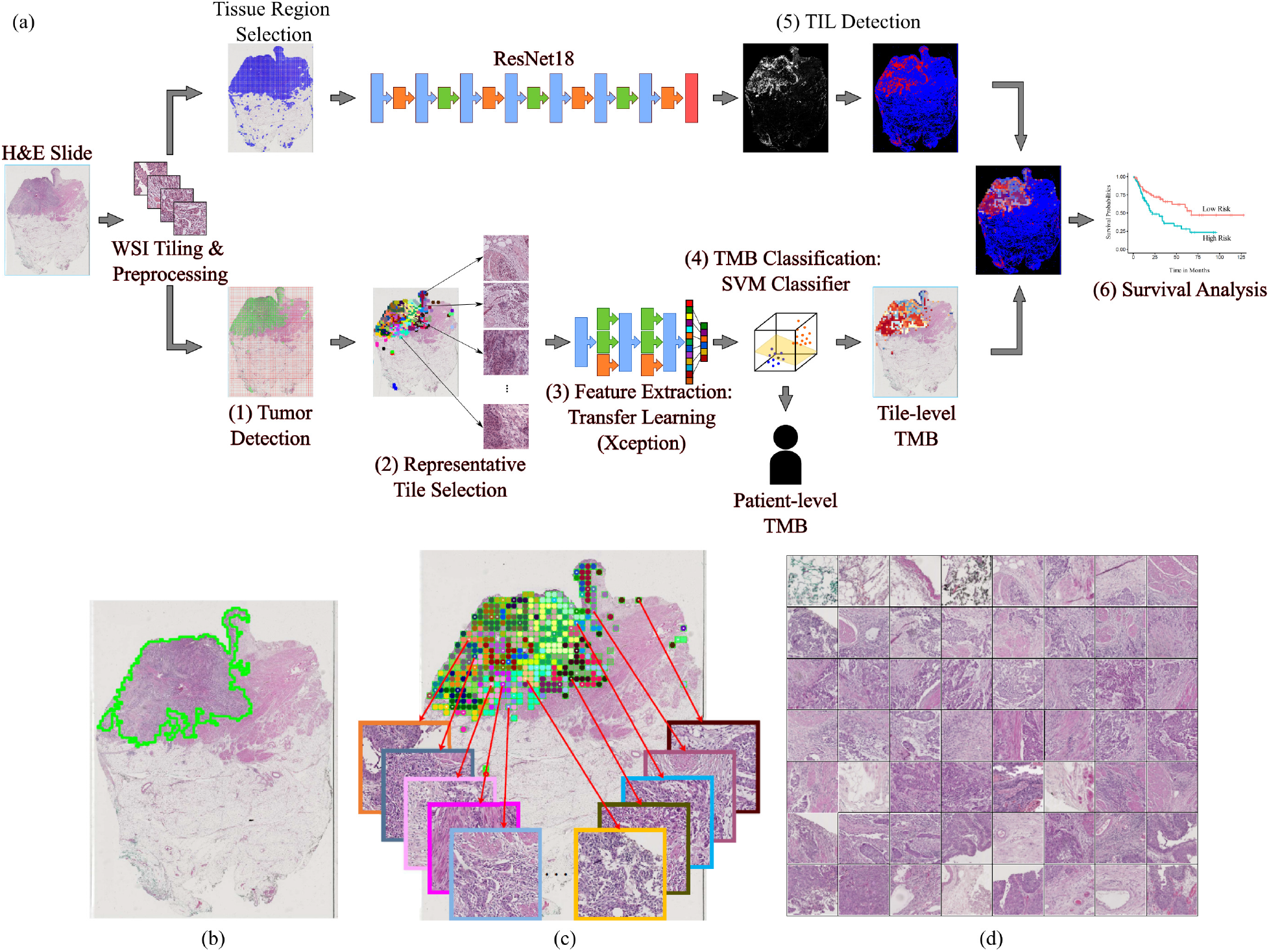
An overview of workflow for our proposed approaches to predict TMB status and TILs from WSIs (a) An Illustration of TMB and TILs pipelines. Given a WSI, we first divide the WSI into small tiles (i.e., regions) and perform preprocessing (e.g., color normalization) within the WSI. To predict patient and tile-level TMB status, we first detect tiles carrying tumors and perform AP clustering to select representative tiles. We use Xception model to extract features from the selected representative tiles, then use SVMs to classify patient and/or tile-level TMB status. In parallel, we use ResNet18 model to detect TILs regions within the WSI. We integrate and perform spatial TMB and TILs analysis to identify patient subgroups with distinct overall survival outcome. (b) Tumor detection result (overlapped greencontours). (c) Example of AP clustering on tumor tiles, where tumor tiles belonging to different clusters are indicated by different color of blocks in the image. Several representative tumor tiles indicated by arrows are zoomed-in for better viewing. (d) 56 representative tumor tiles selected by AP clustering for the slide shown in (c).

### Patient cohorts

Two patient cohorts with digitally scanned WSIs were collected from the TCGA project through the Genomic Data Commons Portal (https:// portal.gdc.cancer.gov/). The TCGA BLCA cohort consists of 386 patients (and corresponding clinical information) with 457 diagnostic H&E stained WSIs. The first diagnostic slide image (i.e., with DX1 suffix) was selected if there are multiple diagnostic slide images available for a patient. Based on the percentile of total number single nucleotide variants [2], 386 TCGA BLCA patients were categorized into 3 groups: 128 low, 128 intermediate and 130 high TMB patients. One high and four low TMB patients were excluded due to severe pen marks on slides, thus 124 low and 129 high TMB patients were used to train and test a model to predict patient-level TMB-H and low status. Based on TMB prediction and TILs detection, the whole cohort of patients with survival information was used for prognosis analysis on patients’ overall survivals. While we focused on BLCA cohort, we also collected TCGA LUAD cohort as an additional dataset to evaluate our proposed computational pipelines. In a similar way with TCGA BLCA cohort, 478 TCGA LUAD patients with 541 diagnostic H&E stained WSIs were collected and divided into TMB high, intermediate and low based on the number of somatic mutations. Due to severe pen marks on slides, 18 low and 4 high TMB LUAD patients were excluded from the image analysis. Finally, 140 low and 157 high TMB patients were used to train and test a model to predict patient-level TMB-H and low status in the LUAD cohort.

### Evaluation on patient-level TMB Prediction

We first investigated whether the use of either tumor detection, representative tile selection, or color normalization as well as different transfer learning models could impact the performance of patient-level TMB prediction. Using TCGA BLCA dataset, we ran patient-level TMB prediction experiments by excluding tumor detection (abbreviated as P-E-TD), no representative tile selection (abbreviated as P-E-RTS), or no color normalization (abbreviated as P-E-CN). We also tested transfer learning on two well-known models, Inception-v3 (abbreviated as P-InceptionV3) [15] and Resnet50 (abbreviated as P-Resnet50) [13], in addition to Xception model (abbreviated as P-Xception), to evaluate whether different transfer learning models could impact patient-level TMB prediction performance. We trained SVM classifiers with linear or RBF kernels to predict patient-level TMB status. The leave-one-out cross validation was employed during testing different configurations. ROC curves of patient-level TMB prediction using different settings in our pipeline are shown in Figs. 2(a) and (b) using SVM with linear kernel (Linear SVM) and SVM with RBF kernel (RBF SVM), respectively (see more details in Table s3). The linear and RBF SVMs with P-Xception and P-E-RTS models achieved overall best AUROC values compared to other methods. While both approaches showed good prediction performance, the P-Xception model used the 11,164 selected representative tiles out of 125,358 tiles, which required significantly less computational time (see computational comparison example in Table s4) compared to the P-E-RTS model. This indicates that the use of AP clustering to select a set of representative tiles from a WSI increases computational efficiency without a significant loss of prediction performance. Therefore we used the AP clustering module in our proposed pipeline for further experiments. The patient-level TMB prediction performance using Xception model (P-Xception) is more accurate than those of Inception-v3 (P-InceptionV3) and Resnet50 (P-Resnet50), thus we used Xception model as the transfer learning algorithm in our pipeline for the rest of experiments.

**Figure 2:**
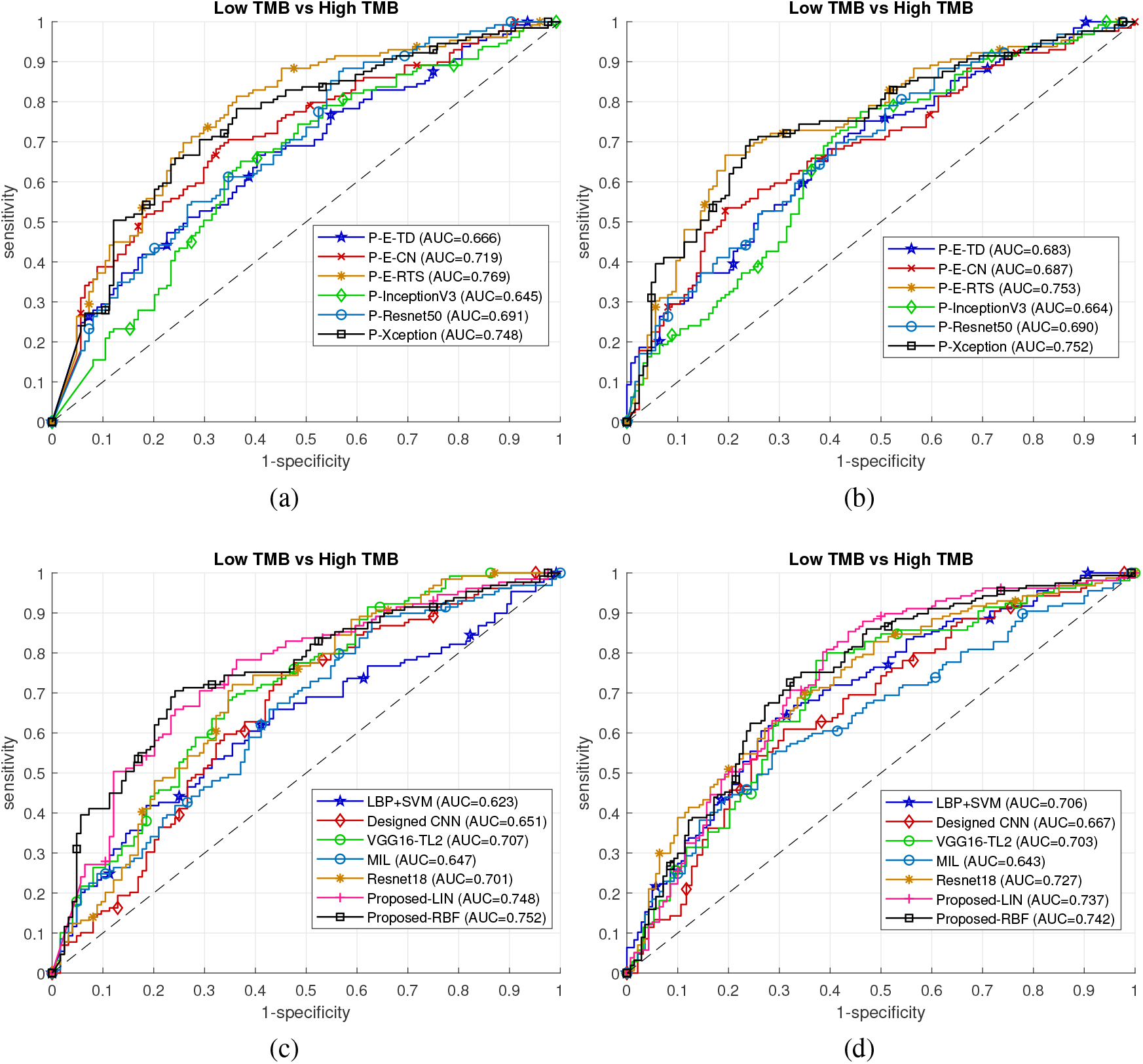
Evaluations on TMB prediction. Ablation study of our method on TCGA BLCA TMB prediction: (a) using SVM with Linear kernel, (b) using SVM with RBF kernel. (c) Baseline comparisons of TCGA BLCA patient-level TMB predictions. (d) Baseline comparisons of TCGA LUAD patient-level TMB predictions. Note that in (c)(d) Proposed-LIN and Proposed-RBF represent the proposed technique using Linear SVM and RBF SVM, respectively

To compare the performance of patient-level TMB prediction with other state-of-the-art methods, we trained our designed CNN model (see Fig.s1 in supplementary methods), VGG16-TL2 [39] and Resnet18 [41], and Multiple Instance Learning based deep learning algorithm [42] as baseline models. To train these deep learning models, tumor tiles of each WSI were assigned the same label (e.g., TMB-H or low status) as the corresponding patient-level TMB status. The final patient-level TMB prediction was obtained by averaging prediction probabilities of all tumor tiles. In addition, we also extracted local binary pattern (LBP) texture features from representative tumor tiles and made predictions using an SVM classifier with RBF kernel as the baseline. Three-fold cross validation was applied to evaluate baseline deep leaning models, due to computational complexity, and the leave-one-out cross validation was used to evaluate the rest methods. Table 1 shows patient-level TMB prediction results in terms of accuracy (*ACC*), specificity (*SPE*), sensitivity (*SEN*) and AUROC values for our proposed method and baseline models. Fig. 2(c) and (d) shows patient-level TMB prediction performance in TCGA BLCA and LUAD, respectively. Overall, the proposed pipeline provides better performance over baseline methods, which achieves from 2% to 5% improvements with respect to AUROC values. Taken together, these results indicate the efficacy of the proposed method to predict patient-level TMB status using WSIs.

**Table 1:**
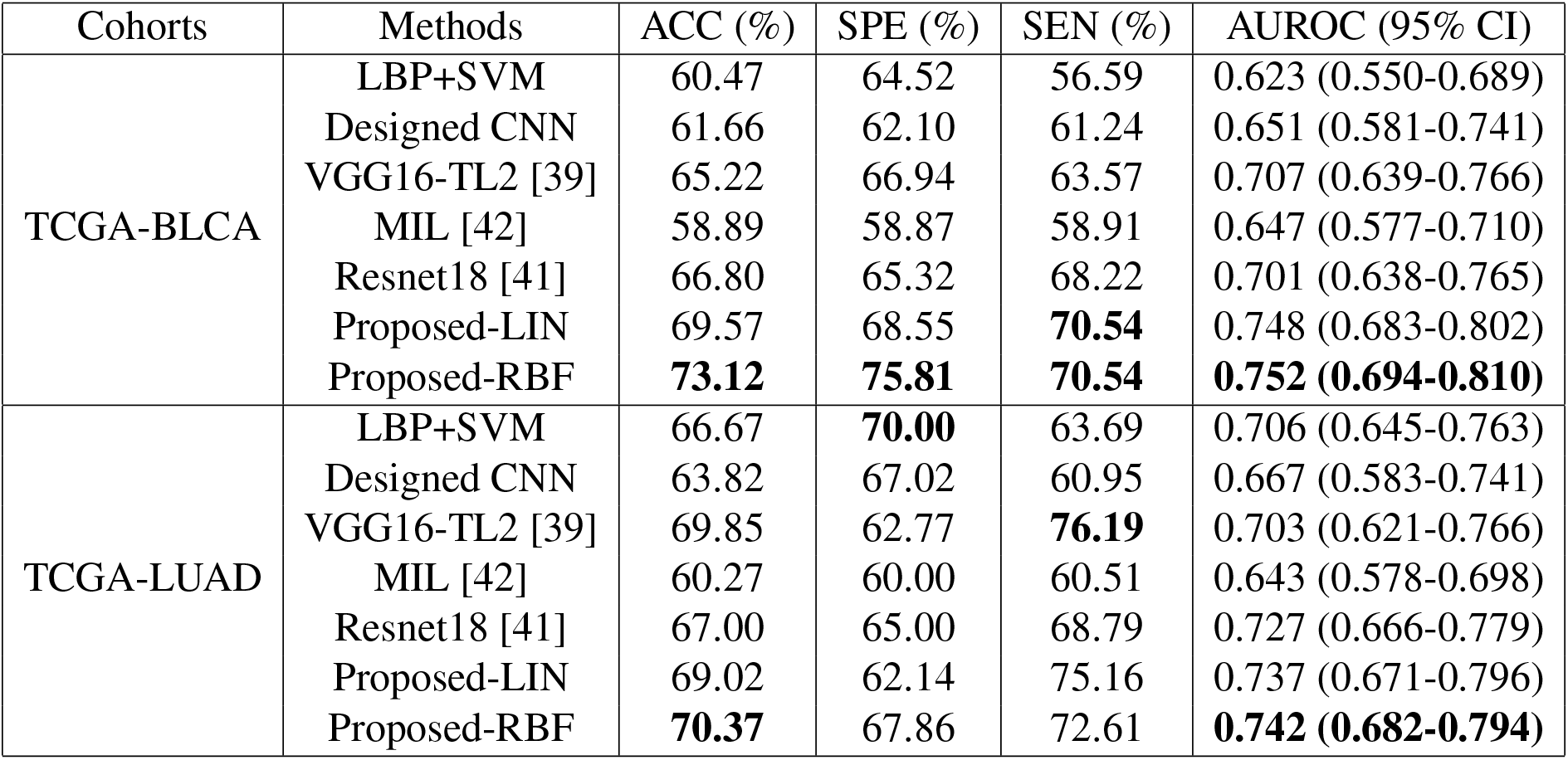
Comparison of patient-level TMB prediction using different methods. In this table, Proposed-LIN uses SVM classifier with linear kernel, while Proposed-RBF uses SVM classifier with RBF kernel.

| Cohorts | Methods | ACC (%) | SPE (%) | SEN (%) | AUROC (95% CI) |
| --- | --- | --- | --- | --- | --- |
| TCGA-BLCA | LBP+SVM | 60.47 | 64.52 | 56.59 | 0.623 (0.550-0.689) |
|  | Designed CNN | 61.66 | 62.10 | 61.24 | 0.651 (0.581-0.741) |
|  | VGG16-TL2 [39] | 65.22 | 66.94 | 63.57 | 0.707 (0.639-0.766) |
|  | MIL [42] | 58.89 | 58.87 | 58.91 | 0.647 (0.577-0.710) |
|  | Resnet18 [41] | 66.80 | 65.32 | 68.22 | 0.701 (0.638-0.765) |
|  | Proposed-LIN | 69.57 | 68.55 | **70.54** | 0.748 (0.683-0.802) |
|  | Proposed-RBF | **73.12** | **75.81** | **70.54** | **0.752 (0.694-0.810)** |
| TCGA-LUAD | LBP+SVM | 66.67 | **70.00** | 63.69 | 0.706 (0.645-0.763) |
|  | Designed CNN | 63.82 | 67.02 | 60.95 | 0.667 (0.583-0.741) |
|  | VGG16-TL2 [39] | 69.85 | 62.77 | **76.19** | 0.703 (0.621-0.766) |
|  | MIL [42] | 60.27 | 60.00 | 60.51 | 0.643 (0.578-0.698) |
|  | Resnet18 [41] | 67.00 | 65.00 | 68.79 | 0.727 (0.666-0.779) |
|  | Proposed-LIN | 69.02 | 62.14 | 75.16 | 0.737 (0.671-0.796) |
|  | Proposed-RBF | **70.37** | 67.86 | 72.61 | **0.742 (0.682-0.794)** |

### Spatial heterogeneity of TMB status correlated with overall survival outcome in BLCA

We investigated whether patient-level TMB status predicted based on WSIs could be useful to identify patient subgroups with distinct clinical outcome on the whole TCGA BLCA cohort. The patient-level TMB status for WES-based TMB high or low group was predicted from our trained SVM with RBF kernel during the leave-one-out cross validation as described in above section. The patient-level TMB status for WES-based TMB intermediate group was independently predicted as TMB high or low by our trained SVM with RBF kernel on WES-based TMB high and low groups. We grouped the whole TCGA BLCA cohort into two subgroups: predicted TMB-High vs TMB-Low, and then generated a Kaplan Meier (KM) plot of the two subgroups using overall survival (OS) (see Fig.s8(a)). While the predicted patient-level TMB-High subgroup shows a trend towards better overall survival (OS), OS difference was not significant between two subgroups using logrank test (P=0.072). We then evaluated if the the spatial heterogeneity of TMB (SH-TMB) within the patient’s tumor could be used to stratify patient subgroups with distinct clinical outcome. We applied the proposed TMB prediction approach on the APC-selected representative tumor tiles. Then, the corresponding tumor regions were assigned the same TMB status of their respective representative tile. To determine the SH-TMB status, we calculated the Shannon entropy [36] of predicted TMB levels of tumor regions within the WSI, i.e., 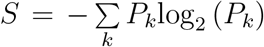, where *P_k_* is the ratio between the number of the *kth* unique TMB prediction probability and the total number of tumor tiles within the WSI. A high entropy value indicates high SH-TMB (e.g., mixture of predicted TMB-H and low regions), while low entropy value indicates low SH-TMB within a tumor (e.g., either TMB-H or low status across most of tumor regions within WSIs). High or low entropy status was determined by using the median entropy value from all patients of TCGA BLCA cohort as the threshold (see Fig.s10(a), Table s10). Fig. 3 shows a visualization of SH-TMB heatmaps based on tile-level TMB prediction, where red and blue colors indicate predicted TMB-H and low status probability, respectively. Fig. 3(a) shows a SH-TMB heatmap of TMB-H patient based on Whole Exome Sequencing (WES) data. Our WSI-based method correctly predicted the patient-level TMB status. The entropy value based on tile-level TMB prediction indicated low SH-TMB. Specifically, the heatmap showed that most tumor regions within the WSI presented TMB-H status, while few tumor regions presented TMB low status. Similarly, Fig. 3(b) showed that our WSI-based method correctly predicted the patient level TMB status as TMB low and low SH-TMB for the WSI-based TMB low patient. Fig. 3(c) and (d) showed that while WSI-based patient level TMB status of these two patients agreed with WES-based patient level TMB status, there are different mixtures of TMB-H and low status within tumor regions. Higher entropy values based on tile-level TMB status indicate higher degree of SH-TMB within WSIs.

**Figure 3:**
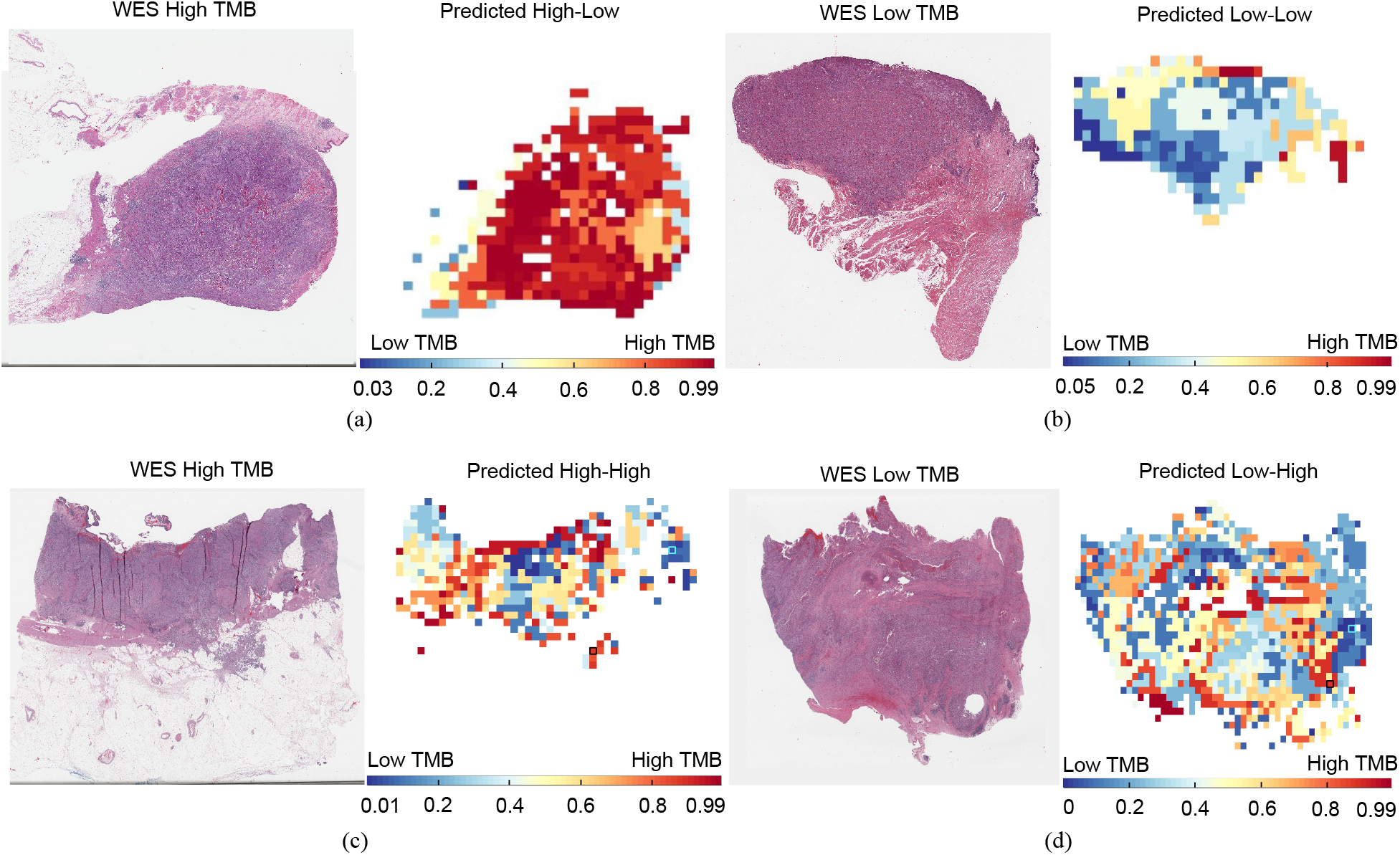
Tile-level TMB prediction visualization. (a) Tissue-based TMB-H patient (TCGA-XF-AAN2) was predicted as patient-level TMB-H based on our WSI-based method. A heatmap of tile-level TMB prediction across tiles (i.e., tumor regions) and entropy measurement showed that most of tumor regions have TMB-H status (i.g., low SH-TMB). (b) Tissue-based TMB low patient (TCGA-XF-A9SH) was predicted as patient-level TMB low and low SH-TMB based on our WSI-based method. (c) Tissue-based TMB-H patient (TCGA-DK-A3IT) was predicted as patient-level TMB-H, while tile-level TMB prediction indicated the high SH-TMB. (d) Tissue-based TMB low patient (TCGA-FD-A3B7) was predicted as patient-level TMB low with high SH-TMB.

To investigate the prognostic utility of SH-TMB status, we selected patient subgroups by utilizing both patient-level TMB prediction and SH-TMB status. In experiments using TCGA BLCA cohort, we predicted patient-level TMB status for 368 patients using our proposed WSI-based method. For each patient, we assigned low or high SH-TMB status based on entropy values derived from tile-level TMB prediction. We assigned patients with predicted patient-level TMB-high and low SH-TMB into one subgroup and the rest of patients to the “Others” subgroup. Then, we generated an OS KM plot segregating by these subgroups (Fig. 4(a)), which indicates that the two subgroups have statistically significantly different OS by using log-rank test (P = 0.016). By univariate analysis using Chi-square test, the TMB subtypes correlated significantly with differences in tumor stage (P = 0.024), but not age (Age>60 vs others, P = 0.872), sex (P = 0.086), lymphovascular invasion (P = 0.064) and inflammatory infiltrate response (P = 0.428) (see Table s5). The patients in patient-level TMB-H with low spatial heterogeneity subgroup had more advanced tumor stage. The TMB subtypes did not significantly correlate with known molecular subtypes determined by Reverse Phase Protein Array (RPPA) (P = 0.761) and mRNA subtypes (P = 0.942) from TCGA BLCA study. Multivariable Cox proportional-hazard analyses of cancer stage and TMB subtypes in relation to the risk of death showed that TMB subtypes remained statistically significantly correlated with survival (see Table s6). A KM analysis of TMB subtypes based on both patient-level TMB and SH-TMB status showed that TMB subtypes with high SH-TMB status have worse OS, regardless of patient-level TMB status (see Fig.s8(b)). We further investigated whether incorporating WSI-based patient-level TMB and spatial heterogeneity with tissue-based TMB testing could improve patient stratification. While WES-based TMB-H patients tend to have better prognosis, we hypothesize that integrating WSI-based patient-level TMB status as well as spatial heterogeneity with WES-based TMB status could further improve patient stratification. To test our hypothesis, we selected 126 WES-based TMB-high patients and divided them into two subgroups: 1) WSI-based patient-level TMB-high and low SH-TMB patient subgroup (HHL) and 2) the rest of WES-based TMB-high patient subgroup (w/o HHL), respectively. Fig. 4(b) showed that WES-based TMB-H & WSI-based patient-level TMB-H and low SH-TMB patient subgroup has better OS compared to the other subgroup (log rank test P = 0.018). Taken together, these results indicate that incorporating WSI-based patient-level TMB status with SH-TMB information could lead to better patient subgroup identification with distinct OS outcome.

**Figure 4:**
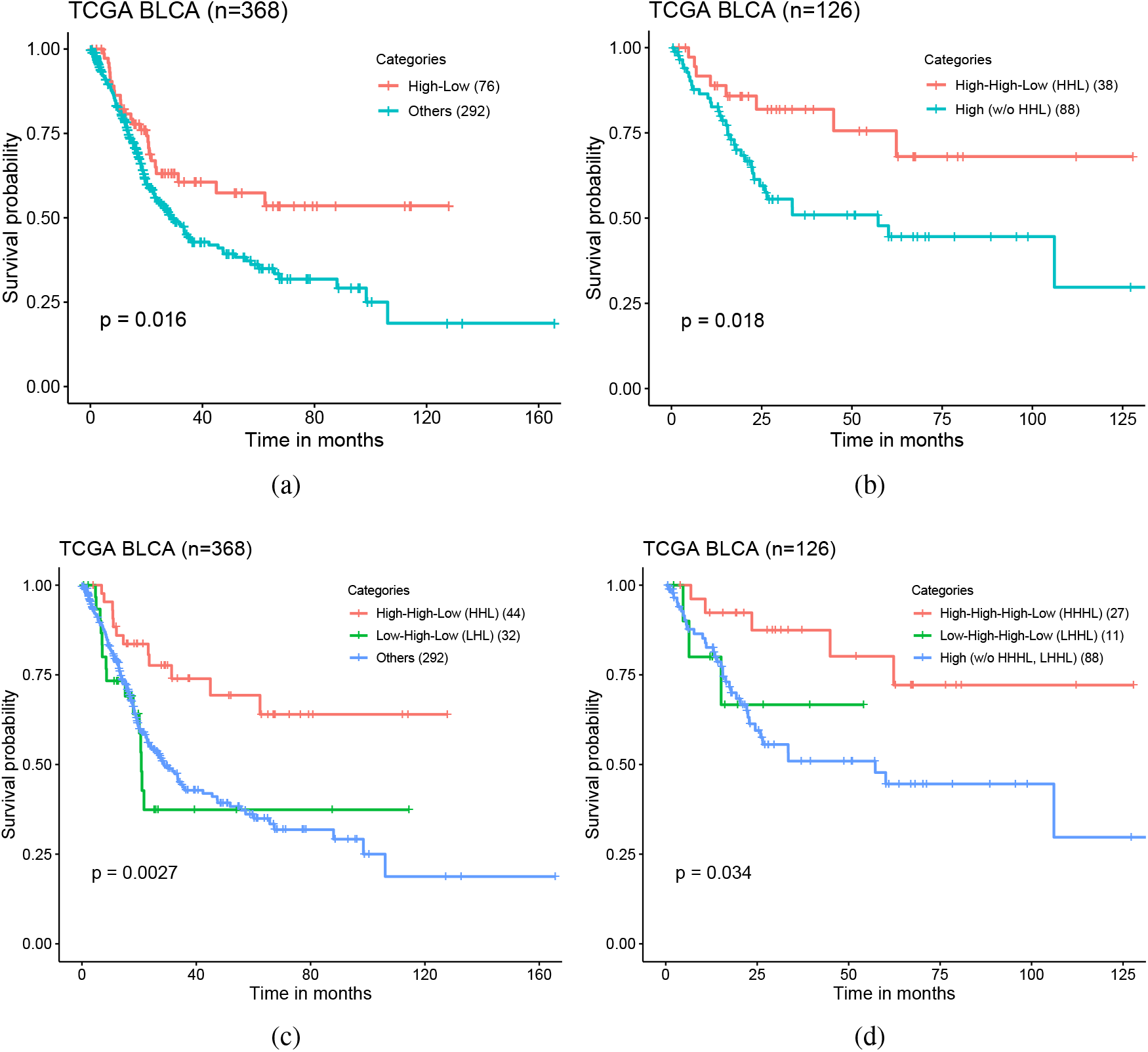
WSI-based patient subtypes. (a) A Kaplan-Meier (KM) plot of overall survival according to WSI-based patient-level TMB-H & low spatial TMB heterogeneity (High-Low) vs other subtypes. (b) A KM plot of overall survival for 126 WES-based TMB-H patients according to WSI-based patient-level TMB-H & low spatial TMB heterogeneity (HHL) vs other WES-based TMB-H subtypes. (c) A KM plot of overall survival according to WSI-based TILs High & patient-level TMB-H & low spatial TMB heterogeneity (HHL) vs WSI-based TILs Low & patient-level TMB-H & low spatial TMB heterogeneity (LHL) vs other subtypes. (d) A KM plot of overall survival for 126 WES-based TMB-H patients according to WSI-based TILs high & TMB-H & low spatial TMB heterogeneity (HHHL) vs WSI-based TILs Low & patient-level TMB-H & low spatial TMB heterogeneity (LHHL) vs other subtypes.

### Spatial analysis of TMB heterogeneity and TILs within tumors further improved patient risk stratification in BLCA

Finally, we investigated that whether the use of spatial co-organization of predicted TMB-H and TILs within the tumor could improve prognostication. We hypothesize that a patient whose majority tumor regions are enriched with TMB-H status (e.g., patient-level TMB-H with low spatial TMB-H heterogeneity) and co-localized with high densities of TILs might have better prognosis. For instance, tumors with patient-level TMB-H and low spatial TMB-H heterogeneity status enriched with high density of TILs (i.e., high number of both TMB-H and TILs regions within the tumor) could show better prognosis compared to patients either having low number of TILs with TMB-H tumors or TMB-L tumors regardless of TILs status. We measured TIL densities within tumor regions for all patients of TCGA BLCA cohort and used the median TIL density score to divide patients into TIL high or low patient subgroups (e.g., >8.12% as TIL high patient subgroup) (see Fig.s10(b)). Then we selected a subset of patients from a TIL high sub-group with the following criteria: predicted TIL **H**igh & predicted TMB **H**igh & predicted **L**ow SH-TMB (HHL). Similarly, to investigate whether high or low level of TILs densities could be linked to patients’ prognosis, we also selected patients from a TIL low subgroup with the following criteria: predicted TIL **L**ow & predicted TMB **H**igh & predicted **L**ow SH-TMB (LHL). Patients belonging to the HHL subgroup tend to have most tumor regions carrying TMB-H status (i.e., a patient-level TMB-H with low SH-TMB) and higher level of TILs co-present within the patient’s tumor (ANOVA testing p<0.001) (see Fig.s11(a)). Fig. 5 shows visualization of TMB-H and TILs carrying regions within the tumors in the HHL, LHL and other subgroups. Fig. 4(c) shows a KM plot of three subgroups (e.g., the HHL subgroup vs the LHL subgroup vs other patients) and a log rank test indicates that three subgroups have statistically significant different OS (P=0.0027). The HHL subgroup showed overall best OS compared to two other subgroups. Multivariable Cox proportional-hazard analyses of cancer stage, lymphovascular invasion, mRNA-based molecular subtype, and joint TIL-TMB based patient subgroups in relation to the risk of death showed that joint TIL-TMB based patient subgroups remained statistically significantly correlated with OS (see Table s7). Interestingly, although patients in the LHL subgroup carry patient-level TMB-H with low spatial TMB-H heterogeneity, the LHL subgroup showed poorer OS compared to the HHL sub-group (HR: 3.30, 95% CI: 1.34-8.12, P<0.01). Lastly, we selected WES-based TMB-H patients and divided into three subgroups: predicted TIL **H**igh & WES-based TMB **H**igh & WSI-based predicted TMB **H**igh & predicted **L**ow SH-TMB (HHHL) vs predicted TIL **L**ow & WES-based TMB **H**igh & WSI-based predicted TMB **H**igh & predicted **L**ow SH-TMB (LHHL) vs other WES-based TMB-H patients. Three subgroups from WES-based TMB-H patients have statistically different TMB-H and TILs overlapped ratio while the HHHL subgroup carrying the highest TMB-H and TILs overlapped ratio among the subgroups (ANOVA testing p=0.005) (see Fig.s11(b)). Fig. 4(d) shows a KM plot of three subgroups and indicates that patients in the HHHL subgroup present better OS than other WES-based TMB-H patient subgroups (log rank test p=0.034). These results show that incorporating TILs density with patient-level and SH-TMB within the tumor based on WSIs and could provide a novel prognostic biomarker to identify high or low risk patient sub-groups.

**Figure 5:**
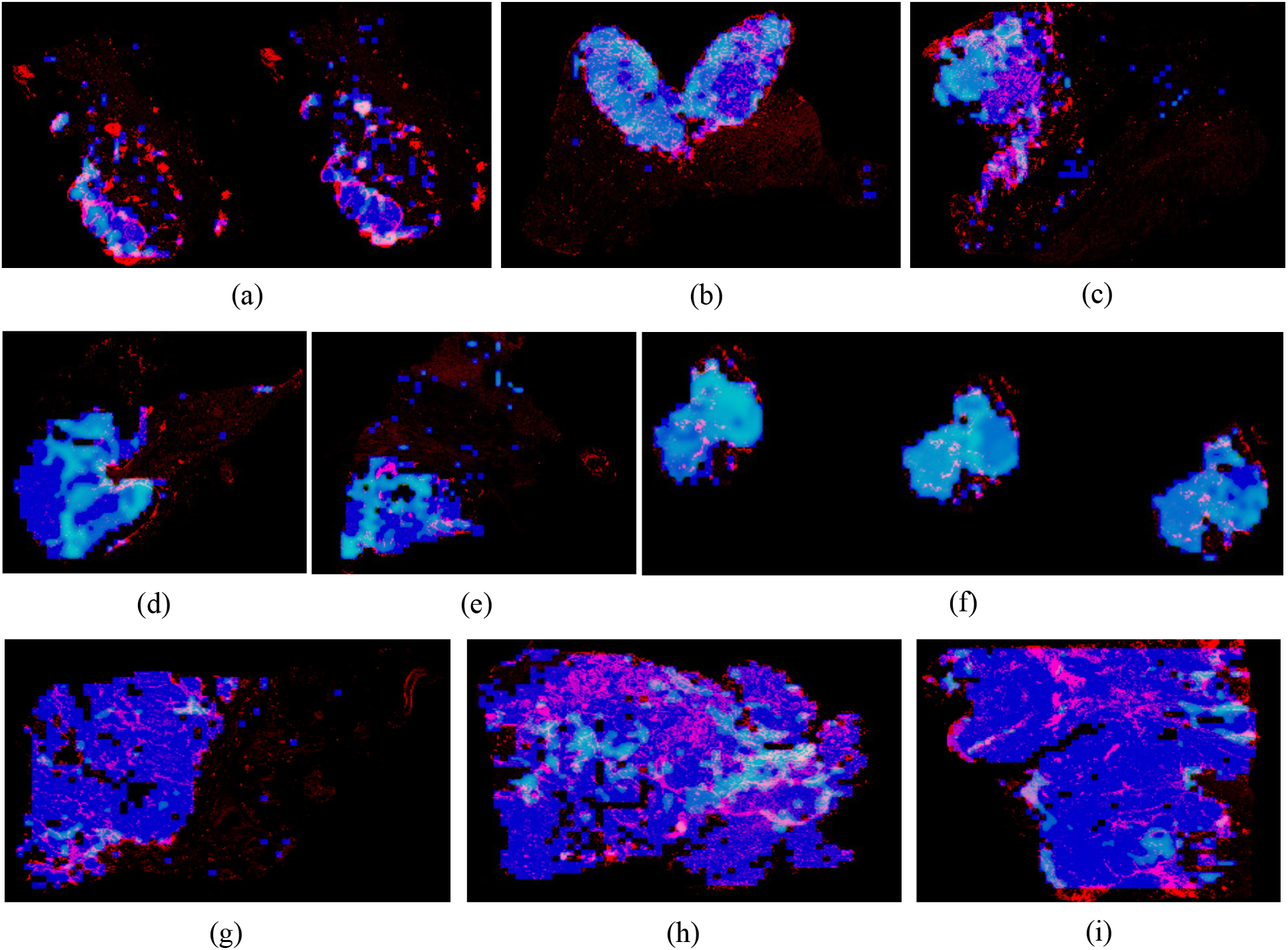
Visualization of spatial heterogeneity and organization of TMB-H and TILs within tumors. Blue color represents identified tissue regions in WSIs. Light blue (e.g., Cyan) color represents predicted TMB-H region. Red color represents predicted TILs. (a)-(c) Patients with high TILs & patient level TMB-H with low spatial TMB-H heterogeneity (e.g., low TMB entropy) patient slides (HHL subtype). Most of tumor regions have been predicted as TMB-H status with high level TILs presence. (d)-(f) Patients with low TILs & patient level TMB-H with low spatial TMB-H heterogeneity patient slides (LHL subtype). Most of tumor regions have been predicted as TMB-H status but with low level TILs presence. (g)-(i) Patients with high TILs & TMB-Low (e.g., others subtype). High TILs present within tumors with predicted TMB-L status.

## 3 Discussion

Intratumor heterogeneity is one of key mechanisms driving disease progression, response and resistance to therapies [29, 31]. Multi-regional tissue-based sequencing from a tumor has shown spatial heterogeneity of mutational signature, mutational burden, T-cell receptor repertoire, etc. [3, 4, 6, 7] and its implication for treatment strategy [8]. While the multi-regional tissue-based sequencing approach could provide landscape of spatial heterogeneity, it is practically challenging to generate such data, due to high costs, limited tissue availability, etc‥ In this study, we present 1) the transfer learning based computational pipeline utilizing WSIs to predict patient-level TMB status and investigate spatial heterogeneity of TMB within tumors. We showed that our proposed computational pipeline could achieve overall best performance to predict patient-level TMB status compared to other state of the art methods. We also showed that measuring and incorporating spatial heterogeneity of TMB status with patient-level TMB status based on WSIs or combined with WES-based TMB status could lead to identify patient subgroups with distinct OS outcomes. Specifically, we found that incorporating SH-TMB information with predicted patient-level TMB status could improve patient risk stratification compared to the use of predicted patient-level TMB status alone (See Fig.s8(a) and (b)) in TCGA BLAC cohort. More specifically, patient-level TMB-H with low SH-TMB status was correlated with better OS. Visual inspection of selected tumor tiles from WSIs by our pathologist indicates that predicted TMB-H representative tumor tiles from patient-level TMB-H WSIs are present with higher densities of TILs, while showing more high grade tumors (see Table s9). This is consistent with the univariate analysis of TMB subtypes showing a higher portion of high grade tumors in patient-level TMB-H and low SH-TMB tumors. Although we observe an enrichment of high graded tumors in this TMB-H subgroup, we reasoned that the higher presence of TILs within the tumors from this subgroup might lead better prognosis. To further investigate whether higher level of TILs with SH-TMB within tumors correlates with patients’ OS, we trained end-to-end deep learning models to detect TILs and quantify TILs density within tumor regions. The predicted TILs density scores were incorporated with SH-TMB information within tumors to identify patient subgroups. The survival analysis of patient subgroups with and without high TILs presence within TMB-H tumors showed that patients carrying TMB-H status within most of tumor regions enriched with high number of TILs have statistically significant better OS. It is worth to note that patient subgroup identification and survival analysis using solely TILs high and low densities information (e.g., TILs high vs low) did not show statistically significant OS difference using log rank test in TCGA BLAD and LUAD cohorts (P=0.32 and P=0.35 in Fig.s8(c) and s9(d), respectively), which indicates the importance of joint spatial TILs and TMB analysis as a prognostic biomarker. It is also worth to note that in TCGA LUAD cohort a patient subgroup carrying patient-level TMB-H and low SH-TMB status with high TILs has statistically significant better overall survival compared to another patient subgroup (log rank test P=0.04 in Fig.s9(e)). However, we did not find meaningful correlation among patient subgroups based on other criteria (e.g., patient-level TMB-H and spatial low heterogeneity of TMB-H status vs others in Fig.s9(a)(b)(c)(d)(f)). This may indicate that the correlation between spatial TMB and TILs patterns linked with OS would be present in a specific type of cancers rather than pan-cancer types, and would need for further investigation across cancer types. Nonetheless, our analysis demonstrated the prognostic utility of spatial TMB and TILs information based on WSIs in BLCA cohort. To the best of our knowledge, this is the first study to predict SH-TMB and investigate prognostic utility of spatial organization of TMB and TILs information for patient stratification in bladder cancer.

There are several limitations and challenges in our study. While we showed overall better performance to predict patient-level TMB status compared with baseline methods, a larger independent cohorts from multiple institutes are needed to validate the performance of the proposed pipeline and its generalizability. Our evaluations indicated that various deep learning-based prediction models, including end-to-end deep learning models, to predict patient-level TMB status did not show superior performance. Larger and more well-annotated WSI datasets would be needed to better train and improve the performance of deep learning-based prediction models (and thus our computational pipeline too, since we employ deep learning-based transfer learning models). Our WSI-based image analysis is performed based on a tile-level not a single cell level without distinguishing certain types of immune cells, and did not take into account specific types of spatial arrangement patterns between regions harboring TMB-H and TILs (e.g., TILs densities within local TMB-H clustered regions). For instance, the single cell level lymphocyte/immune cell detection (e.g., CD4+/CD8+/FOXP3+) and joint spatial analysis of TMB-H tumor cell and/or region and TILs and/or more advanced statistical TMB and TILs spatial modeling [10] could provide higher resolution of TMB-H tumor and immune co-localization within tumor and immune microenvironment (TIME).

In summary, this study demonstrates the feasibility of predicting patient-level TMB status and delineating spatial heterogeneity and organization of TMB and TILs by using computational models based on histological WSIs. Our spatial TMB and TILs analysis shows that patients with more homogeneous TMB-H status across regions within the tumor carrying high density of TILs present better prognosis in bladder cancer. Joint spatial analysis of TILs and TMB within TIME for patients’ tumor provides an unique insight into how immune environment might have an influence on prognosis of patients with TMB-H status. By combining tissue-based TMB-H status with image-based TMB-H/L SH-TMB status could further improve patient stratification in bladder cancer. Taken together, our work provides new foundation of how spatial characterization of tumor (e.g., TMB-H status) and immune environment within the tumor based on WSIs could be used to improve risk stratification in bladder cancer.

## 4 Materials and Methods

### Automatic TMB Prediction

Our designed patient-level TMB prediction includes the following four steps. More implementation details and parameter settings could be referred in the supplementary methods.

#### (1) Tumor Detection

We trained a light-weight convolutional neural network (see the architecture in Fig.s1) model with only about 0.28M trainable parameters to detect tumor regions in the WSI. Given the WSI, it is first divided into non-overlapping tiles (512×512 pixels at 20× magnification). The CNN-based tumor detector then predicts each tile as the probability of belonging to cancer regions. The prediction map corresponding to the WSI is generated by stitching predicted probabilities for all image tiles. An empirical threshold (e.g., 0.5) is applied on the prediction map to obtain tumor regions. Our quantitative evaluations showed that the designed CNN-based tumor detector could provide over 90% dice coefficient in bladder cancer detection and a superior performance than several comparative models (see Fig.s6, s7 and Table s1). Fig. 1(b) illustrates an example of cancer detection on a WSI.

#### (2) Representative Tile Selection

To improve computational efficiency in analyzing large predicted tumor regions, we selected a subset of representative tumor regions for analysis. We first divided predicted tumor regions into a set of non-overlapping tiles (128×128 pixels) at 2.5× magnification. We then characterized each tumor tile by a 42 dimensional feature vector (i.e., 40 multi-scale local binary pattern features [17] and 2D location of the tumor tile). After that, affinity Propagation (AP) clustering [19] was applied to identify tumor regions containing tiles with similar morphological patterns [44]. The AP clustering simultaneously identified a number of *r* local tumor regions and their representative tiles *R_j_*, where 1 ≤ *j* ≤ *r*. Fig. 1(c)(d) illustrates AP clustering of tumor tiles on a WSI, where tumor tiles belonging to different clusters are indicated by different color of blocks in the image. Note that there are 56 (*r* = 56 for this example) representative tiles selected among 490 tumor tiles for the patient slide shown in Fig. 1(c).

#### (3) Feature Extraction

We used transfer learning on pre-trained deep learning models to generate features for selected representative tumor tiles. First, to suppress the influence of color variations, a color deconvolution based method [20] is utilized to normalize tumor tiles into a standard color appearance. Second, transfer learning on pre-trained Xception [27] model was used to extract features from selected tumor tiles. Given an input tumor tile *R_j_* at 20× magnification (1024×1024 pixels), the transfer learning model outputs a high-level feature representation *V_j_* which is a 2048 dimensional vector (see Fig.s4). Finally, the feature vector 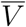 representing the WSI was obtained by integrating features of representative tumor tiles together, i.e., 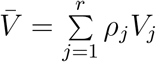, where 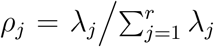 and *λ_j_* represents the number of tumor tiles belonging to the jth cluster. The feature vector 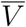 is the weighted mean of features extracted from representative tiles, where each representative tile stands for the major characteristics of tumor tiles within the cluster.

#### (4) TMB classification

We trained the Support Vector Machine (SVM) classifier based on features generated from the transfer learning model to predict patient-level TMB status. First, principal component analysis (PCA) was used to reduce the feature dimension to prevent over-fitting. In this study, we selected the top 100 principal components which provided a superior performance in our testing. Second, feature standardization was performed on each feature component, which ensured its values have zero mean and unit variance. Finally, SVM with radial basis function (RBF) and linear kernels were trained to predict patient-level TMB status.

### TILs Detection

We trained and tested 144 different deep learning models to detect TILs by making use of a public dataset [44], which included 43,440 annotated image tiles. Among 144 trained TIL detectors, the best Resnet18 model provided over the 80% accuracy in distinguishing TIL and Non-TIL tiles during independent testing (see Fig.s5(a) and Table s2), which was selected to perform TIL detection. To identify TIL regions in pathology slides, the WSI was first divided into a set of non-overlapping image tiles (i.e., 112um×112um per image tile). The image tiles were then predicted as TIL tiles or Non-TIL tiles by using the selected TIL detector. The WSI-level TIL detection (see the example shown in Fig.s5(b)) was then generated by stitching tile-level predictions, where tiles with prediction probabilities above 0.5 were considered as TIL regions. Based on tumor and TIL detection, we finally computed the ratio between the number of TIL pixels and the total number of tumor pixels in pathology slides, which was used as a feature variable to quantify TIL densities within tumor regions.

### Code Availability

Our codes for automatic TMB prediction and patient survival analysis are available at: https://github.com/hwanglab/tcga_tmb_prediction. Our codes for automatic TILs detection are available at: https://github.com/hwanglab/TILs_Analysis.

## Acknowledgements

Put acknowledgements here.

## Correspondence

Correspondence and requests for materials should be addressed to Dr.Hwang.

