## Supplementary material for "Spatial heterogeneity and organization of tumor mutation burden and immune infiltrates within tumors based on whole slide images correlated with patient survival in bladder cancer": supplementary_methods_results.pdf

#### 1. Tumor Detection:

To establish the tumor detector, our cooperated pathologist was asked to annotate tumor regions on a set of representative cancer slides. Since the WSI has a huge size (e.g., a few GB/slide), annotated ground truth regions (at 20x magnification) are first divided into a set of non-overlap image tiles, where each tile has 512x512 pixels. We then train a light-weighted CNN-based tumor detector (with only about 0.28M trainable parameters) for identifying bladder cancer regions in the WSI. The architecture of our trained CNN model is shown in Fig.s1, where the input is an image tile and output is the corresponding probability belonging to cancer regions. Our trained CNN model consists of interleaving convolution and pooling layers. Drop-out layers following dense connections are added to against over-fittings. During training, image augmentations including rotation, zooming, flipping and color-based augmentations were randomly applied. During testing, the WSI is first divided into a set of non-overlapping tiles, and then is predicted to be a probability map. An empirical threshold (e.g., 0.5) is applied to binarize tumor regions. Small holes of tumor regions are filled by morphological processing.

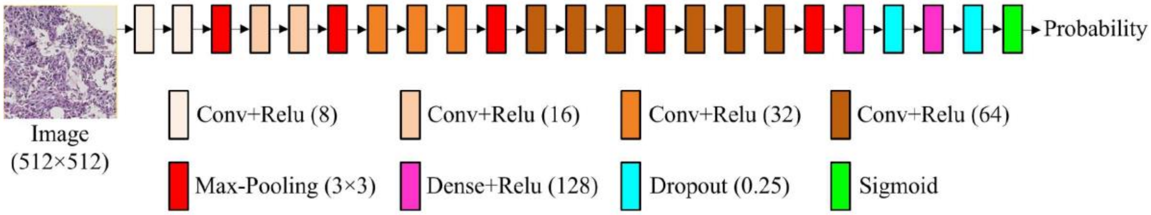


Fig.s1. Architecture of our designed CNN-based tumor detector.

In this study, 100 patients' slides were randomly selected from the whole TCGA BLCA cohort by the pathologist (Dr. Sung Hak Lee), who manually annotated visible tumor regions using ImageScope software from Aperio. Among 100 annotated pathology slides, 48 WSIs (with 64,333 tumor tiles and 73,478 non-tumor tiles) were used for training, while 12 WSIs (with 15,414 tumor tiles and 19,319 non-tumor tiles) were used for validation. The other 40 WSIs were used for independent tumor detection testing. Three evaluation metrics were used, including sensitivity (SEN), precision (PRE) and dice coefficient (DSC). For comparison, we implemented three transfer learning models: VGG16-TL1, VGG16-TL2, and Inception-v3-TL. VGG16-TL1 was trained by fine-tuning three more trainable layers added on top of VGG16 [1], while VGG16-TL2 was trained by fine-tuning the last three convolutional layers [1]. Inception-v3-TL was trained by fine-tuning the last fifteen trainable layers. The number of trainable layers for baseline comparisons was empirically selected based on our training data size (e.g., avoid over-fitting). All models were trained by using RMSprop optimizer to minimize binary cross entropy loss. The maximum training epoch was set as 100. The early stopping was applied if there was no performance improvement on validation set. The batch size was set as 128. The learning rate was set as 0.001. The comparative tumor detection results are listed in Table s1, where 0.5 was used to binarize tumor prediction probability maps. For detail experimental results, please see Table s1, Fig.s6, Fig.s7 in supplementary results.

#### 2. Representative Tile Selection:

Once tumor regions were detected, representative tumor tiles were selected by affinity propagation (AP) clustering [2] for accelerating subsequent feature extraction. The selection of AP clustering rather than other popular clustering techniques such as k-means algorithm is mainly because it does not require to pre-define the number of clusters and provides unique solution with different runs of the algorithm. To efficiently select representative tiles, the detected tumor regions are first divided into a set of 128x128 tumor tiles (at 2.5x magnification). The multi-scale local binary pattern (LBP) texture features are then computed from every image tile, which produces a 40 dimensional feature vector [3]. Note that low resolution tumor tiles are analyzed here, which can help in accelerating computations and capturing macro-scale information from the image by using LBP descriptors. Since geometrically close tumor tiles usually have similar texture patterns, we further incorporate the 2-D location of image tile as the features. Thus each tumor tile is characterized by a 42 dimensional feature vector. The AP clustering then treats each feature vector as a node in a network and recursively transmits real-valued messages along edges of the network until it finds a good set of exemplars and corresponding clusters. As suggested by the reference [2], we define the similarities between feature vectors of tumor tiles as the negative square Euclidean distance between them. The “preferences” that influence the number of finally generated clusters are set as the median value of similarities between feature vectors. Fig.s2 illustrates an example of AP clustering for selecting representative tumor tiles from a whole slide image.

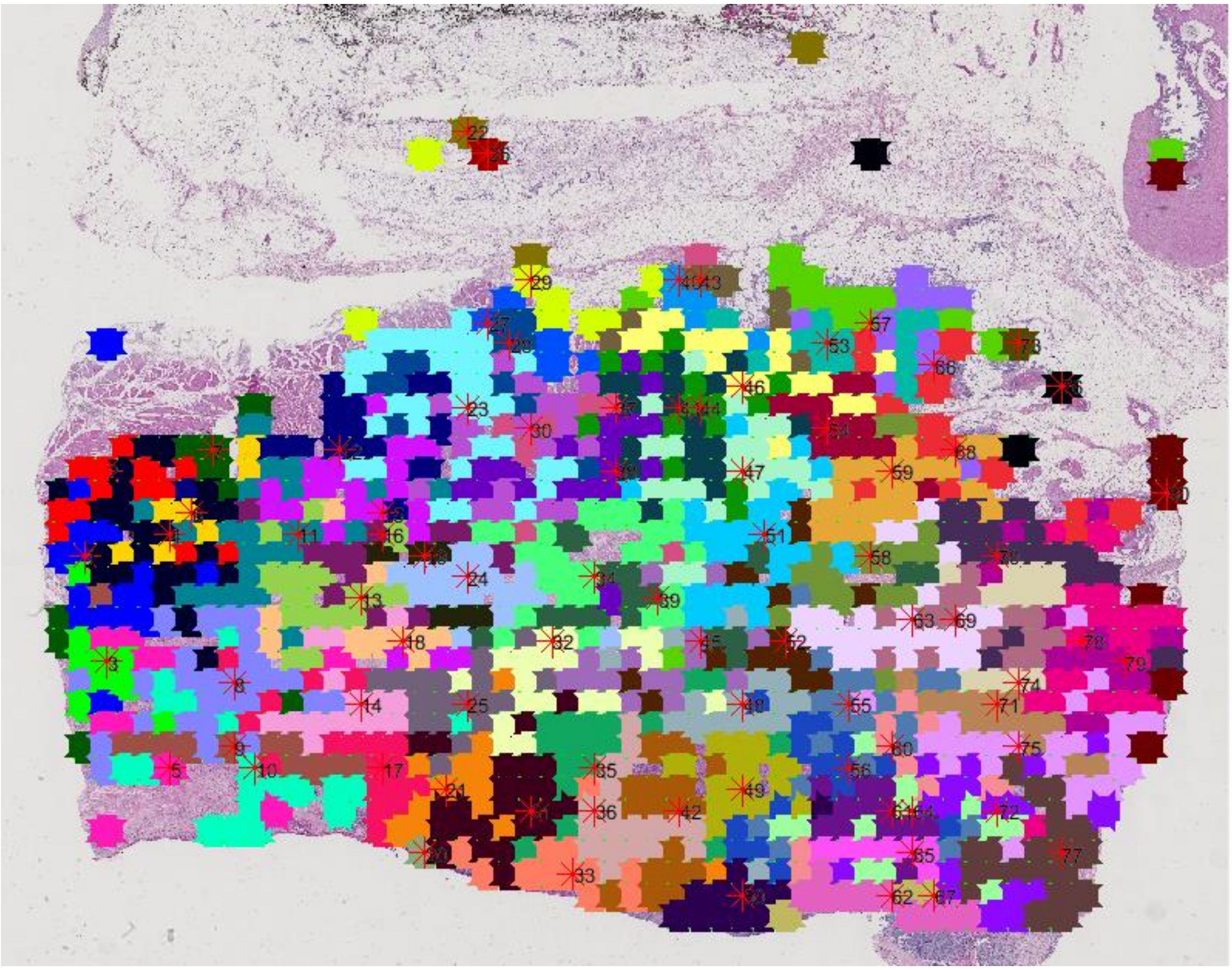

Fig.s2. Example of affinity propagation clustering on a whole slide image. In the image, different color of blocks represents different clusters. One representative tumor tile is selected from each cluster. Note that in this example there are 80 representative tumor tiles which are selected from 1149 tumor tiles. The locations of selected tiles are indicated with red stars and numbers.

#### 3. Feature Extraction:

In this module, we analyze representative tumor tiles and compute a high-level feature representation for the WSI. Since some of TCGA pathology slides only have the highest magnification at 20x, all representative tumor tiles are extracted at 20x for high-level feature extraction. This module includes the following three steps:

(1) Color normalization: Because TCGA pathology images were collected from many different institutions, there exist severe color variations due to different staining procedures. To suppress the influence of color variations, a color deconvolution based method [4] is utilized to normalize tumor tiles into a standard color appearance. Fig.s3. shows an example of color normalization, where the first row shows original tumor tiles and the second row shows the tumor tiles after color normalization.

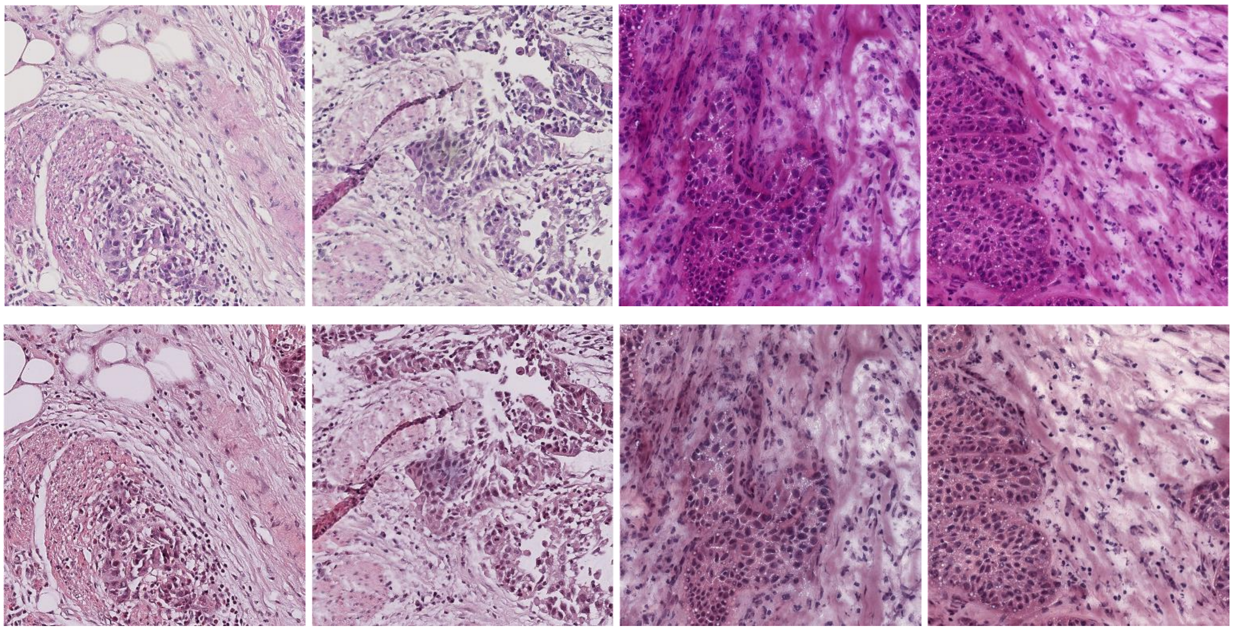

Fig.s3. Illustration of tumor tile color normalization. In the image, the first row shows four examples of tumor tiles before color normalization, while the second row shows correspondingly tumor tiles after color normalization.

(2) Transfer learning: Transfer learning on pre-trained convolutional neural networks have been applied for different classification problems, such as skin cancer [5] and breast cancer [6]. Unlike these transfer learning studies where all images have explicit labels, this study explores to predict TMB from gigabyte whole pathology slides. Tumor representative tiles are determined from WSI for analysis, but they may not contain information relevant to the class assigned to the WSI. In other words, there is no guarantee that a patient slide has the same label with its tumor representative tiles. Therefore, instead of fine-tuning pre-trained models directly for TMB prediction, we make use of pre-trained models as the feature extractor. Motivated by the superior performance on ImageNet classification, we utilize a Xception CNN architecture [7] for transfer learning. The Xception module replaces regular convolutions with depthwise separable convolutions, which has shown an improved classification performance over Google Inception module and residual learning. We remove the output layer of the pre-trained Xception model and re-use all other weights trained by ImageNet. Given an input tumor tile at 20x magnification (with a size of 1024x1024 pixels), the transfer learning model outputs a high-level feature representation which is a 2048 dimensional vector. Fig.s4 illustrates transfer learning using the pre-trained Xception model.

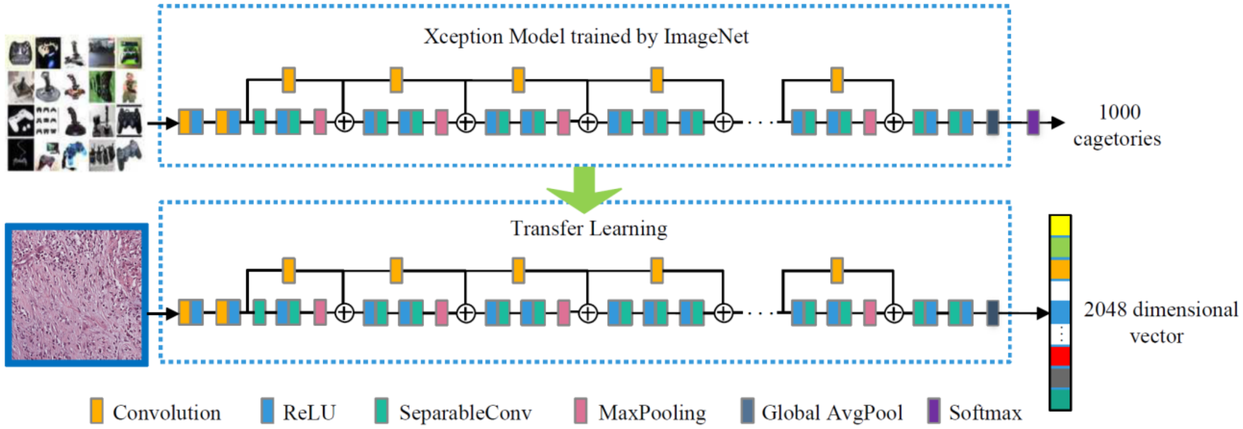

Fig.s4. Illustration of transfer learning on Xception model trained by ImageNet.

(3) Feature integration: After extracting histological features from all representative tumor tiles, the feature vector representing the whole patient slide is obtained by integrating features of individual tumor tiles together. The integration is performed by using the weighted mean of features extracted from representative tiles, where each representative tile stands for the major characteristics of tumor tiles within the cluster. If a cluster includes more tumor tiles, its representative tile is assigned a larger weight such that the WSI could be effectively described by representative tumor tiles.

#### 2. TMB Classification:

The SVM classifier was trained to make patient-level predictions with three steps. (1) Since feature vector of the WSI has a high dimensionality (i.e., 2048) that was much larger than the number of patient samples, feature dimension was reduced to prevent over-fitting to the training dataset. To ensure efficiency and simplicity, principal component analysis (PCA) is utilized, which selects a small number of principal feature components (i.e., 100 for TCGA BLCA, 40 for TCGA LUAD) to train the classifier. The number of principal feature components is determined by optimizing the prediction performance during cross-validations. (2) Feature standardization was performed on each feature component, which ensured its values have zero mean and unit variance. (3) SVM classifiers (e.g., with the RBF kernel or Linear kernel) are trained to predict patient-level TMB status.

After building up the TMB prediction classifier, the testing is performed following a similar procedure of the training process. First, feature dimension reduction and standardization are performed based on the PCA transformation matrix and scaling factors computed from training samples. Then, the trained SVM classifier is applied to predict the patient slide into either low or high TMB category.

#### 3. TILs Detection:

We trained and applied deep learning model to detect TILs regions in WSI. To train the TILs detector, we made use of a public dataset, where 43,440 image tiles were adopted [8]. These image tiles consist of 21,698 TILs patches and 21,742 Non-TILs patches, with dimensions of 150umx150um or 250umx250um. We selected the centering 112umx112um regions from these image tiles for training and testing evaluation. The whole data set was randomly divided into 3 parts: training (80%), testing (10%) and validation (10%). Since TILs detection is a challenging problem, we trained 3 different deep learning models: Resent18, Resent34 and Shufflenet. Image augmentations including random flipping and color jittering were applied along with training. We trained TILs detectors by freezing different percentile of trainable layers, and using different parameter configurations in terms of optimizer, batch size and learning rate. Table s2 lists the grid-search of different parameter settings. Overall, we trained 144 different TIL detectors. Fig. s5 (a) shows testing accuracies of 144 models with different configurations. The best TILs detector which was trained by fine-tuning all trainable layers of Resnet18 and using Adam optimizer with the learning rate of 0.0001 and batch size of 4 provides the best test accuracy (80.06%). The best TILs detector was selected and utilized for TILs detection from the WSI. For example, given a WSI image, the TILs detection was performed by first dividing the WSI into a set of 112umx112um image patches which were predicted as the probabilities belonging to TILs. The WSI-level TILs prediction was finally obtained by stitching tile-level predictions. Fig. s5(b) illustrates an example of TILs detection, where red pixels indicate detected TIL regions in the WSI.

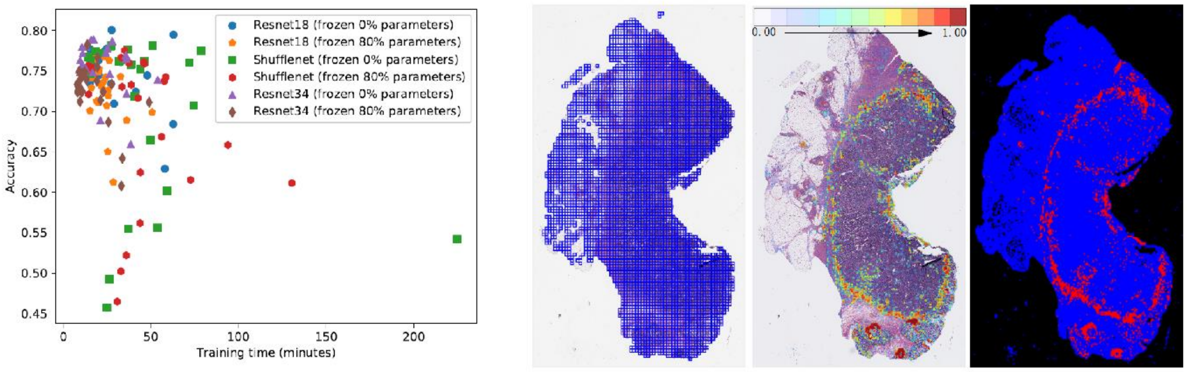


(a) (b)

Fig.s5. TILs detection evaluation. (a) Evaluation of 144 trained TIL detectors. In the figure, the horizontal axis corresponds to the training times of different models, while the vertical axis corresponds to testing accuracies of different models. Note that different symbols (e.g., circles, squares, triangles and pentagons) represent different models with different frozen ratios. Each symbol (e.g., square) has 24 copies, which corresponds to different parameter settings in terms of optimizers, batch sizes and learning rates. It could be found that two Resnet18 models with 0% frozen ratios provide noticeable higher accuracies than other models. (b) Example of TILs detection result on a pathology slide. The first image shows the WSI overlapped with image blocks. The second image shows the WSI overlapped with TILs detection heatmap. The third image shows TILs detection result, where red pixels indicate TILs regions.

### Supplementary Results

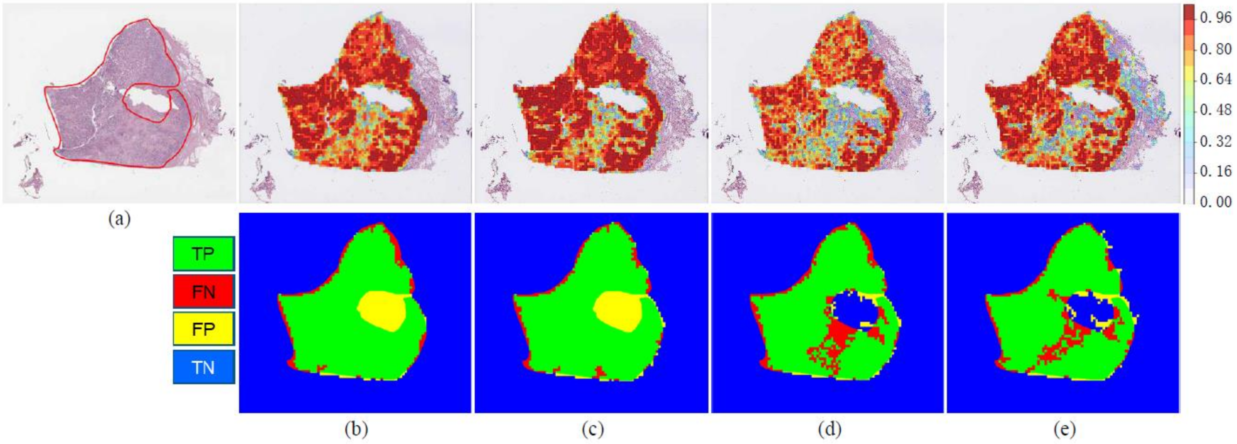


Fig.s6. Example of bladder cancer detection by different models. (a) A WSI with labeled (red) tumor boundaries. (b) VGG16-TL1. (c) VGG16-TL2. (d) Inception-v3-TL. (e) Designed CNN tumor detector. The first row of (b)-(e) shows prediction heatmaps overlapped on WSIs. The second row of (b)-(e) shows predictions comparing with manual annotations. As shown in (e), the designed CNN provides less FP and FN for this example.

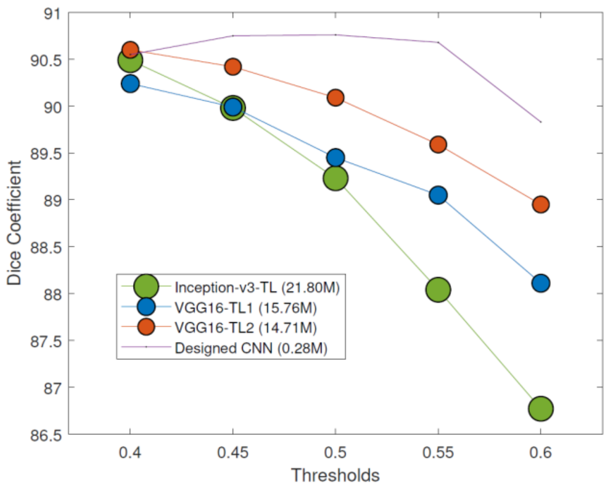


Fig.s7. Dice coefficients of bladder cancer detection with different thresholds, i.e., from 0.4 to 0.6. The size of circles proportionally corresponds to different model size. The designed CNN tumor detector continuously provides higher DSC values than other models. In addition, the designed CNN tumor detector has only about 0.28M trainable parameters, which is more computationally efficient for making predictions on WSIs than other models (with over 10M trainable parameters).

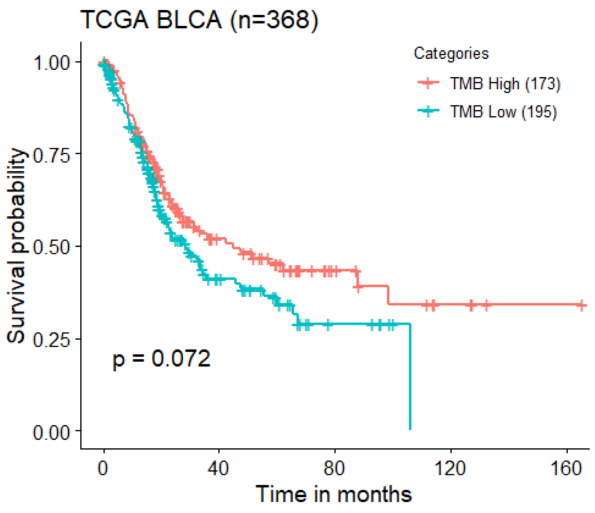


(a)

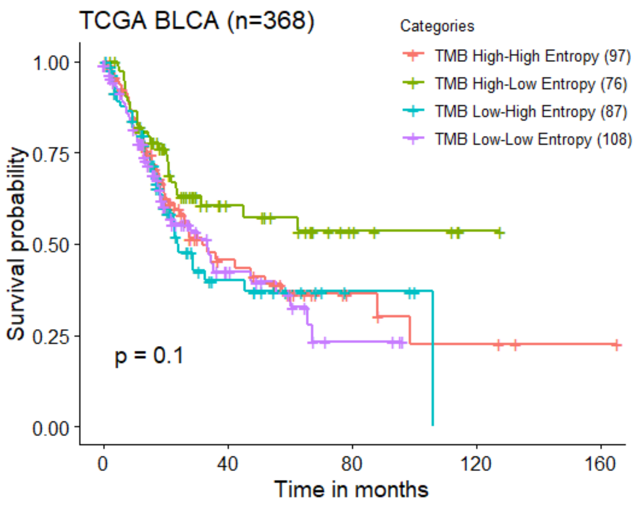


(b)

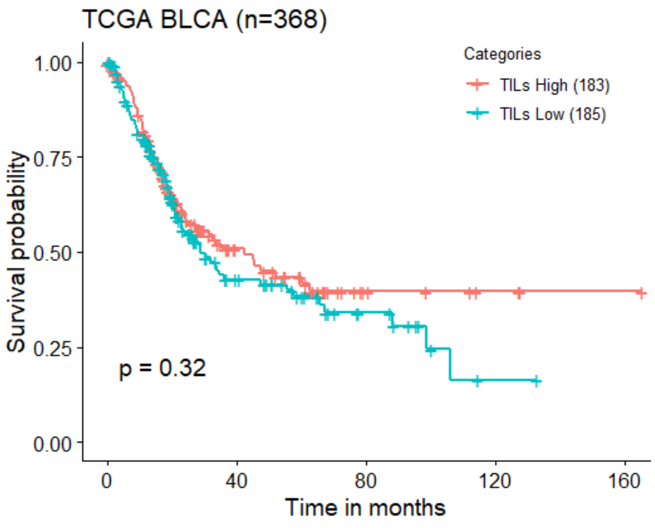


(c)

Fig.s8. KM curves on TCGA bladder cohort. (a) A KM survival curve of TMB subtypes based on predicted patient-level TMB high and low using WSI. Red color represents predicted TMB high subgroup and blue color represents predicted TMB low subgroup (log-rank test P-value = 0.072). (b) A KM survival curve of TMB subtypes based on predicted patient-level TMB status and tile-level entropy analysis (e.g., High-High: WSI-based TMB high and high entropy of tile-level predictions). (c) A KM survival curve based on predicted TILs high and low using WSI. The median level of TILs densities (see Fig. s9(b)) is used as the cut-off value.

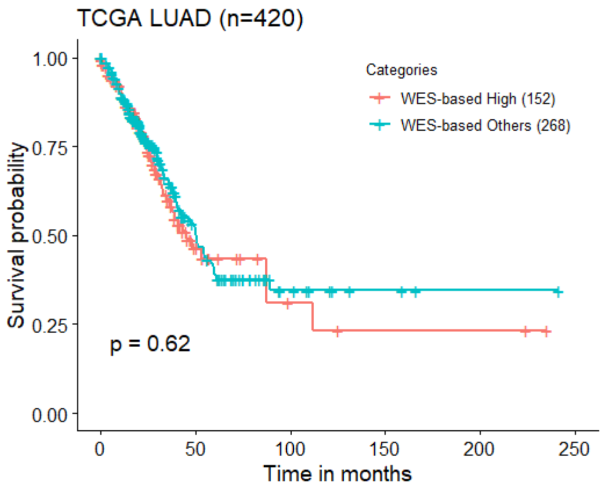

TCGA LUAD (n=420)
Categories
WES-based High (152)
WES-based Others (268)
Survival probability
1.00
0.75
0.50
0.25
0.00
p = 0.62
0
50
100
150
200
250
Time in months


(a)

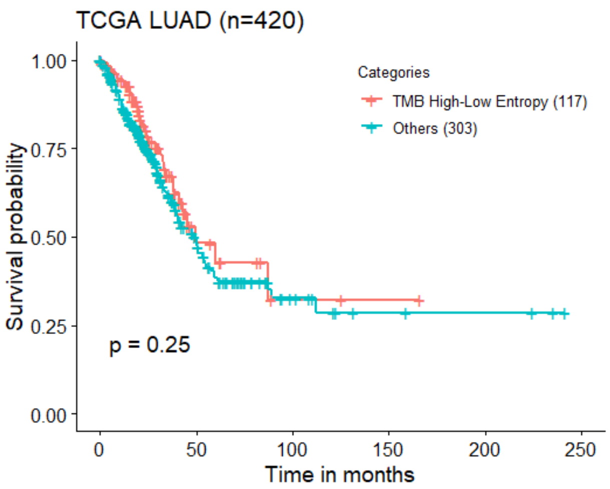

TCGA LUAD (n=420)
Categories
TMB High-Low Entropy (117)
Others (303)
Survival probability
1.00
0.75
0.50
0.25
0.00
p = 0.25
0
50
100
150
200
250
Time in months


(b)

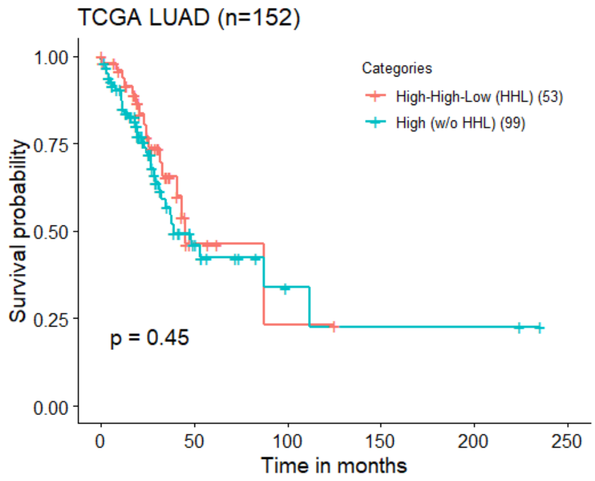

TCGA LUAD (n=152)
Categories
High-High-Low (HHL) (53)
High (w/o HHL) (99)
Survival probability
1.00
0.75
0.50
0.25
0.00
p = 0.45
0
50
100
150
200
250
Time in months


(c)

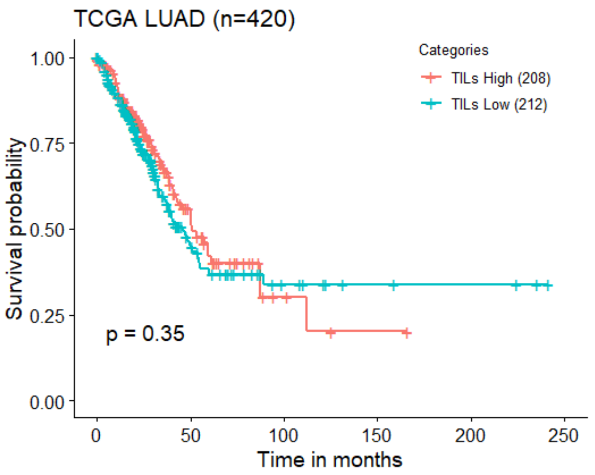

TCGA LUAD (n=420)
Categories
TILs High (208)
TILs Low (212)
Survival probability
1.00
0.75
0.50
0.25
0.00
p = 0.35
0
50
100
150
200
250
Time in months


(d)

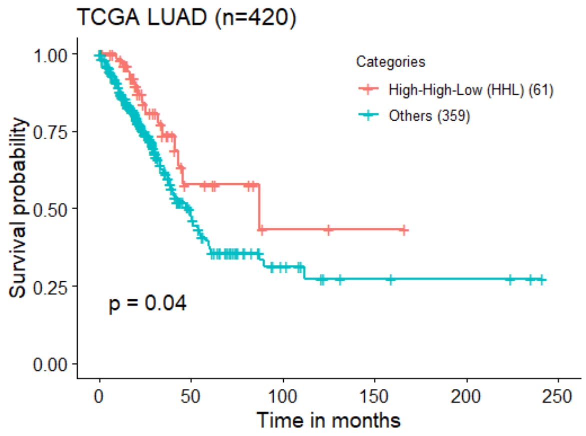

TCGA LUAD (n=420)
Categories
High-High-Low (HHL) (61)
Others (359)
Survival probability
1.00
0.75
0.50
0.25
0.00
p = 0.04
0
50
100
150
200
250
Time in months


(e)

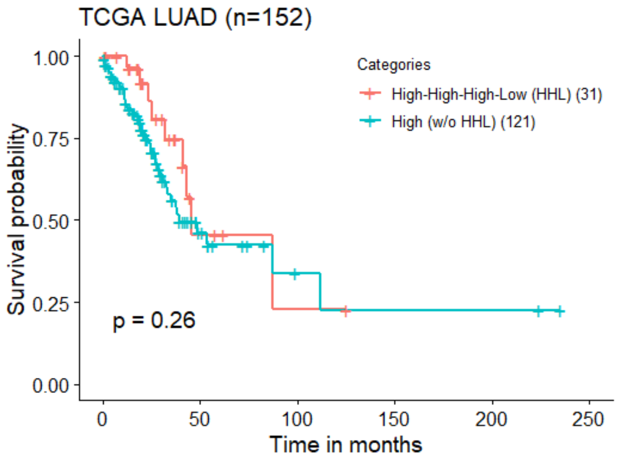

TCGA LUAD (n=152)
Categories
High-High-High-Low (HHL) (31)
High (w/o HHL) (121)
Survival probability
1.00
0.75
0.50
0.25
0.00
p = 0.26
0
50
100
150
200
250
Time in months


(f)

Fig.s9. KM curves on TCGA lung cohort. (a) A Kaplan-Meier plot for overall survival according to WEX-based TMB high & low. (b) WSI-based TMB high & low spatial TMB heterogeneity vs other subtypes. (c) WSI-based TMB high & low spatial TMB heterogeneity vs other subtypes for 152 WES-based TMB high patients. (d) WSI-based TILs High vs TILs Low. (d) WSI-based TILs High & patient-level TMB high & low spatial TMB heterogeneity vs other subtypes. (f) WSI-based TILs high & TMB high & low spatial TMB heterogeneity vs other subtypes for 152 WES-based TMB high patients.

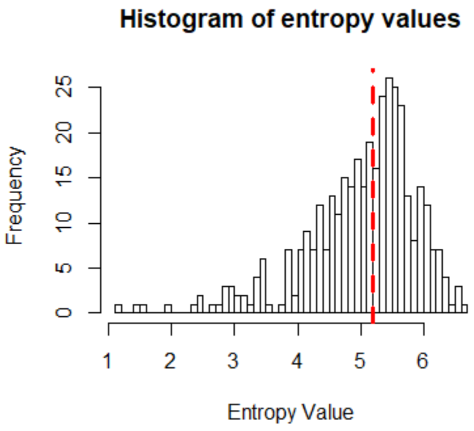


(a)

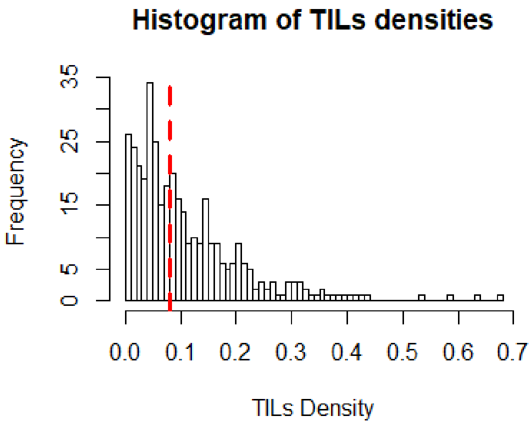


(b)

Fig.s10. Histogram of (a) entropy values of tile-level TMB predictions for TCGA bladder cohort, and (b) TILs densities for TCGA bladder cohort. Note that the red dashed line in (a) indicates the median (5.19) of entropy values on the whole cohort. The red dashed line in (b) indicates the median (0.0812) of TILs densities on the whole cohort.

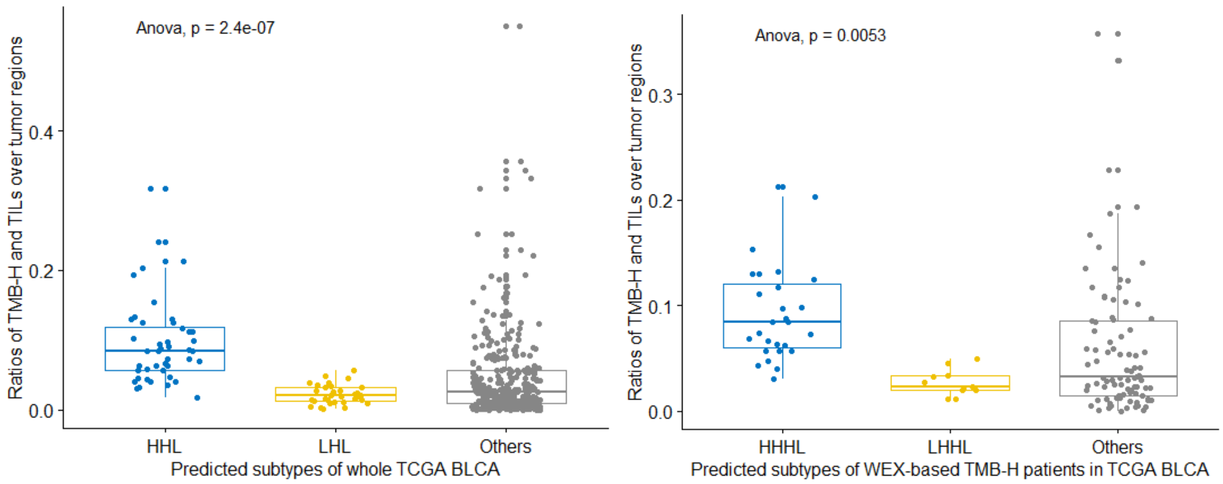


(a) (b)

Fig.s11. Comparison of TMB-H and TILs co-present densities within tumor regions. (a) Three patient subtypes for the whole TCGA BLCA cohort. (b) Three patient subtypes for the WEX-based TMB-H patients in TCGA BLCA cohort.

Supplementary Table s1: TCGA BLCA tumor detection comparative results. Our designed CNN tumor detector provides the highest sensitivity (90.65%) and dice coefficient (90.76%), while Inception-v3-TL provides the highest precision (94.91%).

| Models | SEN (%) | PRE (%) | DSC (%) |
|---|---|---|---|
| VGG16-TL1 | 87.26 | 92.66 | 89.45 |
| VGG16-TL2 | 87.49 | 93.72 | 90.09 |
| Inception-v3-TL | 84.9 | 94.91 | 89.23 |
| Designed CNN | 90.65 | 91.59 | 90.76 |

Supplementary Table s2. Grid Search for finding the best tils detector. By using different parameter settings, 144 different tumor detectors were trained.

| Parameters | Models | Frozen Ratios | Optimizers | Learning Rates | Batch Sizes |
|---|---|---|---|---|---|
| Settings | {'resnet18','resnet34', 'shufflenet'} | {0%, 80%} | {'sgd','adam'} | {'0.001','0.0001','0.00001'} | {4,16,32,64} |

Supplementary Table s3: Ablation study of TMB prediction performance on 253 bladder cancer patients. The values outside and inside the parenthesis are obtained by using SVM classifier with RBF, and Linear kernels, respectively.

| Techniques | Accuracy (%) | Specificity (%) | Sensitivity (%) | AUROC |
|---|---|---|---|---|
| P-E-TD | 64.03(62.06) | 62.90(60.48) | 65.12(63.57) | 0.683(0.666) |
| P-E-CN | 64.43(67.19) | 66.13(69.35) | 62.79(65.12) | 0.687(0.719) |
| P-E-RTS | 71.94(71.15) | 71.77(68.55) | 72.09(73.64) | 0.753(0.769) |
| P-InceptionV3 | 65.22(62.85) | 58.06(59.68) | 72.09(65.89) | 0.664(0.645) |
| P-Resnet50 | 63.64(60.87) | 65.32(54.84) | 62.02(66.67) | 0.690(0.691) |
| P-Xception | 73.12(69.57) | 75.81(68.55) | 70.54(70.54) | 0.752(0.748) |

Supplementary Table s4. Comparison of running time between P-E-RTS (using all detected tumor tiles) and P-Xception (using selected tumor tiles) for the example slide TCGA-2F-A9KO shown in Figure 1(b) of the paper. Our testing was run on a Win10 desktop with Intel(R) i7-7800X CPU, 3.50GHZ, 64GB RAM.

| Methods | Running time (s) |
|---|---|
| P-E-RTS | 2338.20s |
| P-Xception | 402.83s |

Supplementary Table s5. Clinical and pathologic variables of the TCGA BLCA as stratified by the two TMB subtypes.

| | cluster 1 | cluster 2 | total | p-value |
|---|---|---|---|---|
| **N** | 76 (20.7 %) | 292 (79.3 %) | 368 | |
| **Median age (Range)** | 68.5 (45-87) | 69.0 (34-90) | 69.0 (34-90) | 0.902 |
| **Age>60** | 55 (72.4 %) | 214 (73.3 %) | 269 (73.1 %) | 0.872 |
| **Sex** | | | | |
| MALE | 51 (67.1 %) | 224 (76.7 %) | 275 (74.7 %) | 0.086 |
| FEMALE | 25 (32.9 %) | 68 (23.3 %) | 93 (25.3 %) | |
| **Stage** | | | | |
| I | 0 (0.0 %) | 1 (0.3 %) | 1 (0.3 %) | 0.024 |
| II | 28 (36.8 %) | 88 (30.1 %) | 116 (31.5 %) | |
| III | 16 (21.1 %) | 114 (39.0 %) | 130 (35.3 %) | |
| IV | 32 (42.1 %) | 87 (29.8 %) | 119 (32.3 %) | |
| ND | 0 (0.0 %) | 2 (0.7 %) | 2 (0.5 %) | |
| **Lymphovascular.invasion** | | | | |
| NO | 23 (30.3 %) | 95 (32.5 %) | 118 (32.1 %) | 0.064 |
| YES | 36 (47.4 %) | 99 (33.9 %) | 135 (36.7 %) | |
| ND | 17 (22.4 %) | 98 (33.6 %) | 115 (31.3 %) | |
| **Inflammatory.Infiltrate.Response** | | | | |
| ABSENT | 34 (44.7 %) | 116 (39.7 %) | 150 (40.8 %) | 0.428 |
| LYMPHOCYTES | 42 (55.3 %) | 176 (60.3 %) | 218 (59.2 %) | |
| **RPPA.cluster** | | | | |
| 1 | 11 (14.5 %) | 55 (18.8 %) | 66 (17.9 %) | 0.761 |
| 2 | 12 (15.8 %) | 59 (20.2 %) | 71 (19.3 %) | |
| 3 | 13 (17.1 %) | 44 (15.1 %) | 57 (15.5 %) | |
| 4 | 10 (13.2 %) | 36 (12.3 %) | 46 (12.5 %) | |
| 5 | 14 (18.4 %) | 53 (18.2 %) | 67 (18.2 %) | |
| ND | 16 (21.1 %) | 45 (15.4 %) | 61 (16.6 %) | |
| **mRNA.cluster** | | | | |
| BASAL_SQUAMOUS | 27 (35.5 %) | 98 (33.6 %) | 125 (34.0 %) | 0.942 |
| LUMINAL | 5 (6.6 %) | 18 (6.2 %) | 23 (6.3 %) | |
| LUMINAL_INFILTRATED | 17 (22.4 %) | 55 (18.8 %) | 72 (19.6 %) | |
| LUMINAL_PAPILLARY | 22 (28.9 %) | 103 (35.3 %) | 125 (34.0 %) | |
| NEURONAL | 4 (5.3 %) | 15 (5.1 %) | 19 (5.2 %) | |
| ND | 1 (1.3 %) | 3 (1.0 %) | 4 (1.1 %) | |

Supplementary Table s6. Multivariate Cox proportional analysis of tumor stage and the TMB subtypes in relation to the risk of death in the TCGA BLCA cohort.

| | Hazard Ratio (95% CI) | P-value |
|---|---|---|
| Stage* | | |
| III vs II | 1.42287 (0.91257, 2.21852) | 0.11965 |
| IV vs II | 2.95049 (1.95561, 4.45152) | 0 |
| TMB others vs patient level TMB high & Low spatial TMB heterogeneity | 1.79586 (1.18052, 2.73194) | 0.00623 |

*Only two stage I patients

Supplementary Table s7. Clinical and pathologic variables of the TCGA BLCA as stratified by the three TMB subtypes, as shown in Fig.5(c) in the paper.

| **Gender** | Hazard Ratio (95% CI) | P-value |
|---|---|---|
| FEMALE vs MALE | 1.39379 (0.90777, 2.14002) | 0.1291 |
| **Age** | | |
| 60 older vs 60≤ | 2.85441 (1.57398, 5.17645) | 0.00055 |
| **AJCC.pathologic.tumor.stage** | | |
| III vs II | 1.18750 (0.61789, 2.28221) | 0.60615 |
| IV vs II | 1.86517 (0.95066, 3.65944) | 0.06986 |
| **Lymphovascular.invasion** | | |
| YES vs No | 2.05379 (1.27877, 3.29854) | 0.00291 |
| **mRNA.cluster** (vs Luminal_papillary) | | |
| Basal_squamous | 1.44447 (0.84275, 2.47582) | 0.18101 |
| Luminal | 1.64039 (0.72750, 3.69880) | 0.23284 |
| Luminal_infiltrated | 1.39544 (0.76291, 2.55242) | 0.27944 |
| Neuronal | 2.67671 (1.10942, 6.45812) | 0.02845 |
| **TMB subtypes** (Fig.5(c)) | | |
| LHL vs HHL | 3.30484 (1.34371, 8.12820) | 0.00923 |
| Others vs HHL | 3.49581 (1.72801, 7.07214) | 0.0005 |

Supplementary Table s8. Clinical and pathologic variables of the TCGA BLCA as stratified by the three TMB subtypes, as shown in Fig.5(d) in the paper.

| | | |
|---|---|---|
| **Gender** | | |
| FEMALE vs MALE | 0.60296 (0.15890, 2.28794) | 0.45715 |
| **Age** | | |
| 60 older vs 60⩽ | 7.15549 (2.06602, 24.78246) | 0.0019 |
| **AJCC.pathologic.tumor.stage** | | |
| III vs II | 0.34773 (0.10859, 1.11345) | 0.07524 |
| IV vs II | 0.56182 (0.17744, 1.77884) | 0.32684 |
| **Lymphovascular.invasion** | | |
| YES vs No | 4.63173 (1.65690, 12.94760) | 0.00347 |
| **mRNA.cluster** (vs Luminal_papillary) | | |
| Basal_squamous | 0.80302 (0.30896, 2.08717) | 0.65261 |
| Luminal | 2.58872 (0.61467, 10.90264) | 0.19478 |
| Luminal_infiltrated | 1.31437 (0.43356, 3.98464) | 0.62904 |
| Neuronal | 1.14643 (0.21351, 6.15572) | 0.87339 |
| **TMB subtypes** (Fig.5(d)) | | |
| LHHL vs HHHL | 3.33936 (0.51640, 21.59446) | 0.20549 |
| Others vs HHHL | 4.60703 (1.34988, 15.72338) | 0.01473 |
